# LGR5+ Intestinal Stem Cells Display Sex Dependent Radiosensitivity

**DOI:** 10.1101/2023.12.05.570158

**Authors:** Ryan C. Zitter, Rishi Man Chugh, Payel Bhanja, Subhrajit Saha

**Author notes:** Corresponding Author: Subhrajit Saha PhD Associate Professor, Director of Basic Science Research Department of Radiation Oncology Department of Cancer Biology, The University of Kansas Medical Center MS 4033, 3901 Rainbow Boulevard Kansas City, Kansas 66160. Equally Contributed (Ryan C. Zitter, Rishi Man Chugh, Payel Bhanja).

## Abstract

Radiosensitivity, the susceptibility of cells to ionizing radiation, plays a critical role in understanding the effects of radiation therapy and exposure on tissue health and regeneration. Identifying characteristics that predict how a patient may respond to radiotherapy enables clinicians to maximize the therapeutic window. Limited clinical data suggested a difference in male and female radiotherapy outcomes. Radiotherapy for gastrointestinal malignancy is still a challenge due to intestinal sensitivity to radiation toxicity. In this manuscript, we demonstrated sex-specific differences in intestinal epithelial radiosensitivity. In mice models of abdominal irradiation, we observed a significant increase in oxidative stress and injury in males compared to females. Lgr5+ve intestinal stem cells from male mice showed higher sensitivity to radiation-induced toxicity. However, sex-specific differences in intestinal radiosensitivity are not dependent on sex hormones as we demonstrated similar sex-specific radiosensitivity differences in pediatric mice. In an ex-vivo study, we found that human patient-derived intestinal organoids (PID) derived from males showed higher sensitivity to irradiation compared to females as evidenced by loss of budding crypt, organoid size, and membrane integrity. Transcriptomic analysis of human Lgr5+ intestinal stem cells suggested radiation induced upregulation of mitochondrial oxidative metabolism in males compared to females’ possible mechanism for radiosensitivity differences.

## Introduction

Even as cancer treatment progresses, radiotherapy continues to play a role in nearly half of cancer patients treatments (1). While incredible technological advancements have taken place in radiotherapy, nearly all have been focused on anatomical precision. While this has done much in limiting radiation exposure to normal tissue, empiric dosing and fractionation is a cornerstone of the specialty. Personalization of radiotherapy remains for the most part at the hands of clinical judgment. Tailoring dose to individual patients is critically important. Pauses in radiotherapy caused by radiotoxicity can be associated with a decrease in local control by as much as 16% (2). Identifying patients who are inherently radiosensitive and more prone to these toxicities could receive extended fractionation allowing for better chances of toxicity. Inversely, radioresistant individuals may receive a higher dose, giving them a better chance of local control. Increasing our understanding of an individual’s therapeutic index maximizes our chances of disease control.

The simplest personalization a patient could receive in their therapy design is on the basis of their sex. While poorly understood, what clinical data we do have points to a male and female difference in radiation response. Following the Chernobyl disaster, the male to female birth ratio increased in children born to irradiated men (3). Data mining has also shown that women were at increased risk for endocrine imbalance as well as for thyroid and brain tumors (4, 5). In following survivors of the atomic bombs it has been seen that risk of solid cancers increases at 35% per Gy for men, but by 58% per Gy for women (6). Sex based differences in radiotoxicity have been found clinically in hepatic (7) and thyroid (8) tissues. Preclinical models showing differences in normal tissue radiotoxicity are also described (9, 10). Identifying radiosensitive patients is most important when considering acute radiation effect on the intestines. Notoriously among the most radiosensitive of tissues, the intestinal epithelium is of acute concern when irradiating the abdomen (11). Nearly all patients receiving abdominal radiotherapy will experience some form of gastrointestinal symptoms (12, 13). This is due to high radiosensitivity of intestinal epithelium (14) consisting intestinal stem cell (ISCs) with high self-renewal rate. The intestinal microvasculature has also been proposed as a reason for this heightened sensitivity (15), it has been largely refuted (16) and leaves intestinal stem cells ISCs as a primary suspect for determining factor of intestinal radiosensitivity.

The intestinal epithelium is a single cell layer thick and organized into repetitive crypt-villous units, with stem cells residing at the base of the crypt and renewing cells moving upward in the crypt in a conveyer belt fashion (17). Within the ISCs that reside at the crypt base, several subpopulations exist. Homeostasis of the intestine free from external insult are maintained by crypt base stem cells (CBCs) marked by leucine rich repeat containing G protein coupled receptor 5 (LGR5+) (18). Loss of the LGR5+ stem cells in the radiation response is catastrophic to the architecture of the epithelium, highlighting their essential role in radiation toxicity (19). In addition to these, a population of reserve stem cells (RSCs) reside at the “+4” position, 4 cells above the LGR5+ cells (20). These regenerative stem cells have a greater number of markers. Recent studies have investigated several markers of these +4 RSCs and find that they replenish LGR5+ cells as well as other differentiated cell types upon radiation insult (20, 21).

While there appear to be two distinct populations of stem cells—CBCs and RSCs—CBCs have been found to express RSC markers (22) and that RSC are dispensable in the intestinal radiation response (19). It has also been found that YAP1 signaling induces a pro-survival LGR5+ survival type (23), suggesting that plasticity of the LGR5+ is what is crucial in repopulation. Though it has been found that YAP-1 also leads to transient expansion of RSC (20). Our understanding of whether distinct ISC’s populations or plasticity is the defining aspect of inherent radiosensitivity is still in need of clarification.

Considering this uncertainty, the LGR5+ ISC’s remain our most certain candidate for determination of inherent intestinal radiosensitivity. Therefore, determining a basis for any difference observed in the male and female in the intestinal response lies in the LGR5+ population. In studying the LGR5+ response in this setting, it is impossible to separate this cell population from a discussion of global hormonal differences between the sexes. The pro-growth hormone estrogen is often the cited difference for any apparent radio-resistance observed in the clinic (24-26) and has been described in the laboratory (27). However, recent evidence clearly suggests that sex hormone (28) has no influence in ISC homeostasis and repair. In this study, we attempt to determine if there are significant differences in the intestinal radiation response between males and females. We have demonstrated sex specific differences in LGR5+ ISC radiosensitivity having distinct transcriptomic profile.

## Materials & Methods

### Animals

Eight to 12-week-old male and female C57BL6/J mice, Lgr5-eGFP-IRES-CreERT2 mice, Gt (ROSA)26Sortm4(ACTB-tdTomato-EGFP) Luo/J mice, and B6. Cg-Gt (ROSA)26Sortm9(CAG- tdTomato) Hze/J mice were purchased from Jackson laboratories. Lgr5-eGFP-IRES-CreERT2 mice were crossed with Gt (ROSA)26Sortm4(ACTB-tdTomato-EGFP) Luo/J mice (Jackson Laboratories) to generate Lgr5/eGFP-IRES-Cre-ERT2; R26-ACTB-tdTomato-EGFP (29, 30). In Gt (ROSA)26Sortm4(ACTB- tdTomato-EGFP) Luo/J mice tdTomato is constitutively expressed (independent of Cre recombination) in the membrane of all the cells, and therefore allows better visualization of cellular morphology. Lgr5-eGFP-IRES-CreERT2 mice were crossed with B6. Cg-Gt (ROSA)26Sortm9(CAG-tdTomato) Hze/J mice (Jackson Laboratories) to generate the Lgr5-eGFP-IRES-CreERT2; Rosa26-CAG-tdTomato heterozygote for lineage tracing experiments. All the mice were maintained ad libitum, and all the studies were performed under the guidelines and protocols of the Institutional Animal Care and Use Committee of the University of Kansas Medical Center. All the animal experimental protocols were approved by the Institutional Animal Care and Use Committee of the University of Kansas Medical Center (ACUP number 2019-2487).

### In-vitro Culture of Intestinal Crypt Organoids

Small intestine from male and female C57BL6/J, Lgr5-eGFP-IRES-CreERT2; R26-ACTB-tdTomato-EGFP mice, and non-malignant colon tissue from human surgical specimens was used for Crypt isolation and development of ex vivo organoid culture (31, 32). Human colon tissue samples were sourced from the University of Kansas Medical Center Biospecimen Repository, identified with the reference number HUS#5929. Tissue samples were thoroughly cleaned with cold PBS followed by the scraping and chopping of tissue into 5 mm pieces, followed by thorough washing with cold PBS and incubated at room temperature in a gentle dissociation buffer (Stem cell technologies, Cat No. 100- 0485) for 20-30 minutes with gentle shaking. For mouse tissue, the dissociation buffer was subsequently replaced with cold 0.1% BSA in PBS with 4 times and vigorously pipetted followed by passing through the 70μm filter to get 1 to 4 fractions of the supernatants enriched in crypt stem cells. Human tissue required an additional step of pipetting in ice-cold DMEM with 1% BSA to liberate crypts from the tissue. The mouse and human tissue isolated crypts were centrifuged, resuspended in a small volume of PBS containing 0.1% BSA and were embedded in Matrigel (Corning, Cat No. 356231) and cultured in respective intestinal organoid Medium (Stemcell Technologies) with their respective supplements and gentamycin (50 mg/mL). The quantification of organoids per well wad performed by using EVOS Microscope (Thermo Fisher Scientific). Images of organoids were acquired using the fluorescent (Nikon, 80i) and confocal (Nikon, A1RMP) microscopy. These microscopy imaging enable to both quantify and investigate the structural and cellular characteristics of the organoids, contributing to a comprehensive analysis of their properties and development.

### Irradiation procedure

Abdominal irradiation (AIR) was performed on mice anesthetized with 87.5 mg/kg of Ketamine and 12.5 mg/kg of Xylazine using a small animal radiation research platform (XENX; XStrahl) (0.67 mm Cu HVL) at a dose rate of 2.26 Gy/min at 220 kVp and 13 mA. A 3-cm2 area of the mice containing the gastrointestinal tract (GI) was irradiated thus shielding the upper thorax, head, and neck, as well as the lower and upper extremities, and protecting a significant portion of the bone marrow, to predominantly inducing radiation induced gastrointestinal syndrome (RIGS). For partial body irradiation (PBI), the mouse was irradiated at a dose rate of 2.82 Gy/min at 220 kVp and 13 mA, by shielding one leg from radiation exposure. In order to achieve a uniform distribution of the radiation dose, a deliberate approach was taken in the radiation delivery process. Specifically, half of the total irradiation dose was administered from the anterior-posterior (AP) direction, and the remaining half was delivered from the posterior-anterior (PA) direction. This strategy helps to mitigate the potential for dose inhomogeneity within the target tissue. The total irradiation time required to administer the intended radiation dose was calculated with respect to dose rate, radiation field size, and fractional depth dose to ensure accurate radiation dosimetry.

### Histology

Exposure to radiation doses exceeding 8Gy has been observed to prompt a cascade of cellular responses within the intestinal tissue. Specifically, within a day after irradiation, there is a notable induction of cell cycle arrest and apoptosis among the crypt epithelial cells. These cellular events contribute to a substantial reduction in the population of regenerating crypts within the intestinal tissue, a phenomenon observed by day 3.5 post-radiation exposure. The collective impact of these cellular responses ends in villi denudation by day 7 following radiation exposure (33). Animals were euthanized when moribund or at 3.5 days after AIR for time-course experiments, and intestines were collected for histological analysis. The intestinal tissues of experimental animals underwent a thorough wash in PBS to eliminate intestinal contents. Subsequently, the jejunum was fixed using 10% neutral-buffered formalin followed by embedding in paraffin and sectioning into 5-μm-thick sections for hematoxylin and eosin (H&E) and immunohistochemistry (IHC) staining. All the H&E staining was performed at the Pathology Core Facility at the University of Kansas Cancer Center (Kansas City, KS). The histopathological analysis was conducted for the assessment of crypt proliferation rate, villi length and crypt depth to analyze the structural and cellular alterations within the intestinal tissue morphology.

### Crypt Proliferation Rate

To investigate the villous cell proliferation, jejunum part of the intestinal tissue, was processed for paraffin embedding and Ki67 immunohistochemistry. The tissue sections were processed for deparaffinization, rehydration through a series of graded alcohol solutions, and an overnight incubation at room temperature with a monoclonal anti-Ki67 antibody (M7240 mib1; Dako). Nuclear staining was performed using a chromogenic substrate. streptavidin-peroxidase and diaminobenzidine (DAB). In addition, a light counterstaining with hematoxylin was performed for optimal visualization. The identification of murine crypts within the tissue sections was employed using an established histological criterion as previously reported by Potten et al (34). To quantify the proliferation rate of the cells, the percentage of Ki67-positive cells were counted over the total cell count within each crypt. A total of 50 crypts were counted for each animal.

### Determination of Villi Length and Crypt Depth

The villi length and crypt depth were independently and objectively analyzed and quantitated in a double blinded manner from coded digital photographs from H&E stained slides using ImageJ 1.37 software. The crypt depth was measured in pixels from the bottom of the crypt to the crypt villus junction. Villi length was determined by measuring the length from the crypt villus junction to the villous tip. The measurement in pixels was converted to length or depth (in μm) by dividing with the conversion factor 1.46 pixels/μm.

### qPCR analysis

qPCR was performed to determine the oxidative stress responsive genes, mitochondrial biogenesis genes, and stem cells marker and Wnt target genes expression in the male and female intestinal epithelial cells. Total RNA was isolated from the samples using RNeasy micro/mini kit from (Qiagen, Germantown, MD, USA). The concentration and purity of the extracted RNA were checked using a NanoDrop spectrometer (Thermo Scientific, Waltham, MA, USA). 1 μg of total RNA was reverse transcribed using RNA to cDNA EcoDry™ Premix (Double Primed) (Takara Bio USA Inc., San Jose, CA, USA) according to the manufacturer instruction. qPCR was performed using the QuantStudio™ 7 Flex Real Time PCR System (Applied Biosystems™, New York, NY, USA) and SYBR Green Supermix (Bio-Rad, Hercules, CA, USA) with specific primers to the target genes in a 20 μL final reaction volume. GAPDH was used as a reference gene for sample normalization. The primer sequences are listed in Supplementary Table 1. The delta-delta threshold cycle (ΔΔCt) method was used to calculate the fold change expression level in the samples.

### Western blot analysis

Total OXPHOS (oxidative phosphorylation) expression was analyzed in irradiated male and female isolated intestinal epithelial cells. Isolated cells were lysed with 1 × RIPA buffer (Cell Signaling, MA, USA) containing a protease and phosphatase inhibitor cocktail (Thermo Fisher Scientific Inc.) to isolate total protein. The total protein concentration of the samples was determined by the BCA method. For analysis, 30 µg of protein samples with 1 × gel loading buffer were separated by SDS-PAGE and transferred to PVDF membranes. After protein transfer, the membrane was blocked with 5% skim milk for one hour at room temperature followed by overnight incubation at 40C with total OXPHOS antibody cocktail (Abcam, ab110413). After incubation with primary and secondary antibodies, the membrane was developed with Trident Femto Western HRP substrate (GeneTex, Irvine, CA, USA) and imaged using a ChemiDoc XRS + molecular imager (Bio-Rad). β-Actin was used as an internal control for normalization.

### In-vivo Lineage Tracing Assay

To investigate the role of Lgr5-expressing intestinal stem cells (ISCs) in tissue regeneration under homeostatic conditions and in response to irradiation injury an in-vivo lineage tracing assay was performed. Lgr5-eGFP-IRES-CreERT2 mice were crossed with B6. Cg-Gt (ROSA)26Sortm9(CAG-tdTomato) Hze/J mice (Jackson Laboratories) to generate the Lgr5-eGFP- IRES-CreERT2; Rosa26-CAG-tdTomato heterozygote mice. To initiate lineage tracing, adult mice were injected with tamoxifen (Sigma; 9 mg per 40 g body weight, intraperitoneally) in Cre reporter mice, allowing for the labeling of Lgr5+ lineages and their subsequent tdTomato (tdT)-positive progeny. This approach enabled the identification and tracking of Lgr5 ISC lineages. For irradiation injury studies, mice were given 12.5Gy of AIR, and tissue was harvested on day 10 post-irradiation.

### LGR5+ Cell Isolation

Human tissues from, males and females were received from the University of Kansas Medical Center Biorepository (HSC #5929) and dissociated into single cell suspension using gentle MACS dissociator (Miltenyi Biotec, Inc., CA, USA) and maintained at 4°C throughout preparation. Cell numbers were determined using an automated Cell Counter T20 (Bio-Rad). Cell suspension was then centrifuged at 300xg for 10 minutes and supernatant was aspirated. A buffer solution composed of PBS with 0.5% bovine serum albumin (BSA), and 2 mM EDTA was then added at 70 µl per 107 cells per cell pellet. Subsequently, 10 µl of FcR blocking reagent was added per 107 of cells and 20 µl per 107 cells of Anti-LGR5 MicroBeads (Miltenyi Biotec, Inc., CA, USA) were added to the final cells suspension and incubated for 20 minutes in dark at 4°C. Cells were then washed by adding 2 ml of the above buffer and centrifuged at 300xg for 10 minutes. Cells pellets were then resuspended in 500 µl of buffer and passed through a MACS magnetic column while in a magnetic field in order to capture cells labeled with MicroBeads. After allowing cells suspension to run through magnetic column, place the column in a collection tube. The magnetically labelled cells were flushed out by firmly pushing the plunger into the column and rinsed with buffer in order to collect tagged cells in collection tube.

### RNAseq analysis

Isolated human LGR5+ cells were stored at −80°C until processed for RNA isolation using RNeasy micro kit as per manufacturer instruction (Qiagen, Germantown, MD, USA). RNA processing, cDNA generation, mRNA-Seq library prep, and library quality control were performed at the UNC Advanced Analytics Core. RNA was cleaned and concentrated using the Zymo RNA Clean & Concentrator kit (Zymo Cat # R1013). Full-length cDNA was generated using the TaKaRa SMART-Seq v4 Ultra Low kit (TaKaRa Cat # 634889). mRNA-Seq libraries were then prepared using the Illumina Nextera XT DNA Library Preparation kit (Illumina Cat # FC-131-1024). 1 ng of cDNA was used as input and 12 amplification cycles were used during PCR enrichment. Prepared libraries were quantified using the Qubit dsDNA Quantitation High Sensitivity kit (ThermoFisher Cat # Q32851), and fragment size was assessed using the Agilent Bioanalyzer High Sensitivity DNA kit (Agilent Cat # 5067-4626). Libraries were then pooled in an equimolar fashion. The library pool was sequenced at the UNC CRISPR Screening Facility using the Illumina NextSeq 500 instrument. A High-Output 75-cycle flow cell (Illumina Cat # 20024906) was used for single-end sequencing with a read length of 76 bp, PhiX concentration of 1%, and loading concentration of 1.65 pM. This sequencing process ensures the acquisition of high-quality data for downstream analysis. The Resulting gene expression levels were transformed using a variance stabilizing transformation (VST) in order to compare normalized levels across all the tested samples. Heat maps were created using the online software Morpheus from the Broad Institute in addition to running a student’s t-test to compare male and female. Resulting mRNA with p value ≤0.05 underwent gene enrichment using Gene Ontology Enrichment Analysis with the PANTHER classification system (35).

### Statistical analysis

Mouse intestinal sampling regions were chosen at random for digital acquisition for quantitation. Digital image data was evaluated in a blinded manner as to treatment. A two-sided Student’s t-test was used to determine significant differences between experimental groups. All data are presented as mean ± standard deviation (SD). A difference between groups with ∗p<0.05, ∗∗p<0.005, or ∗∗∗p<0.0005 was considered statistically significant.

## Results

### Female Intestinal Epithelium is resistant to radiation induced toxicity

In this study, our aim was to investigate potential differences in radiosensitivity between male and female within the context of the intestinal epithelium. Specifically, we wanted to discover whether there exists an inherent sex-based variation in how these tissues respond to radiation exposure. Additionally, we aimed to ascertain whether such differences, if present, could be attributed to differences in the LGR5+ cell populations between males and females. As LGR5+ cells are commonly associated with intestinal stem cells (ISCs). These cells play a pivotal role in maintaining the regenerative capacity of the intestinal lining, where constant cell turnover is essential for tissue homeostasis.

Histopathological analysis of the intestinal tissue (36) after three days post 12.5 Gy AIR (Figure 1A) demonstrated significantly increased in villi length (p<0.0005) (Figure 1B, C) and crypt depth (p<0.0005) (Figure 1B, D) in adult female C57BL/6 mice compared to male. Higher number of Ki-67 positive proliferating cells per crypt was observed in female intestinal epithelium (p<0.0005) (Figure 1B, E) compared to male.

**Figure 1:**
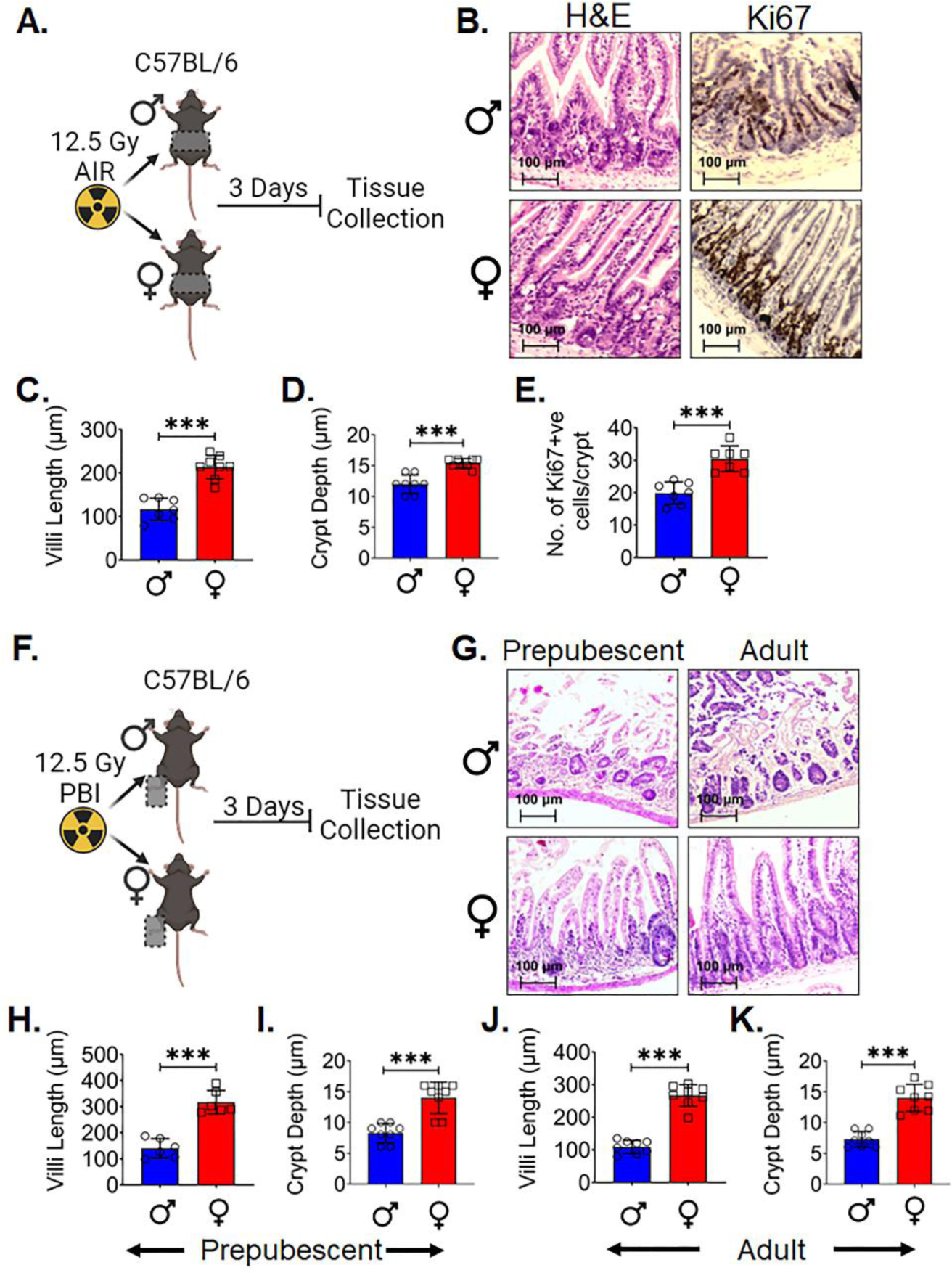
Female Intestinal Epithelium is resistant to radiation induced toxicity: A. Schematic representation of the experimental design, B. Histological analysis of intestinal tissue. Adult female C57BL/6 mice exhibited significantly increased C. villi length and D. crypt depth compared to males, along with a E. higher number of Ki-67 positive proliferating cells per crypt. F. Schematic representation of the experimental design using prepubescent (4 weeks old) and postpubescent (12 weeks old) male and female mice to investigate the role of systemic hormonal differences. G-K Histological analysis of intestinal tissue of both age groups showed that female had significantly increased villi length and crypt depth in comparison to males. Data presented as the mean ± SD. (Significant level, ***: p<0.0005).

In order to determine if finding was dependent on systemic hormonal differences such as estrogen, A separate experiment was performed in both pre (4 weeks old) and postpubescent (12 week old) mice. Male and female C57BL/6 mice 4 week and 12 week old was exposed to lethal dose of 12.5Gy partial body irradiation (PBI) keeping one hind leg out of radiation field for partial bone-marrow sparing. Post 3 days of irradiation tissue were collected for histological analysis (Figure 1F). H&E staining of both the age group mice shows that female mice have significantly increase in villi length (p<0.0005 & p<0.0005 respectively) and crypt depth (p<0.0005 & p<0.0005 respectively) in comparison to the male mice (Figure 1G-K). These data indicates that differences observed in radiosensitivity are independent of major hormonal differences between male and female mice.

### LGR5+ ISCs in female are more radio-resistant compared to male

To examine the radiosensitivity of ISCs we determined the survival of LGR5+ cells in Lgr5/eGFP-IRES-Cre-ERT2; R26-ACTB-tdTomato-EGFP mice. Age matched male and female mice were irradiated with 12.5Gy of abdominal irradiation (AIR) (Figure 2A). As the regeneration of the intestinal epithelium follows the apoptotic phase at 2-3 days (37), assessing the LGR5+ population 3 days post irradiation will provide an accurate depiction of the surviving fraction. Histological analysis of post 3 days of irradiation shows a significant difference in the number of crypts containing LGR5+ cells in female mice compared to the male mice (p<0.05) (Figure 2B, C). This suggests that female LGR5+ cells are more radioresistant than male mice.

**Figure 2:**
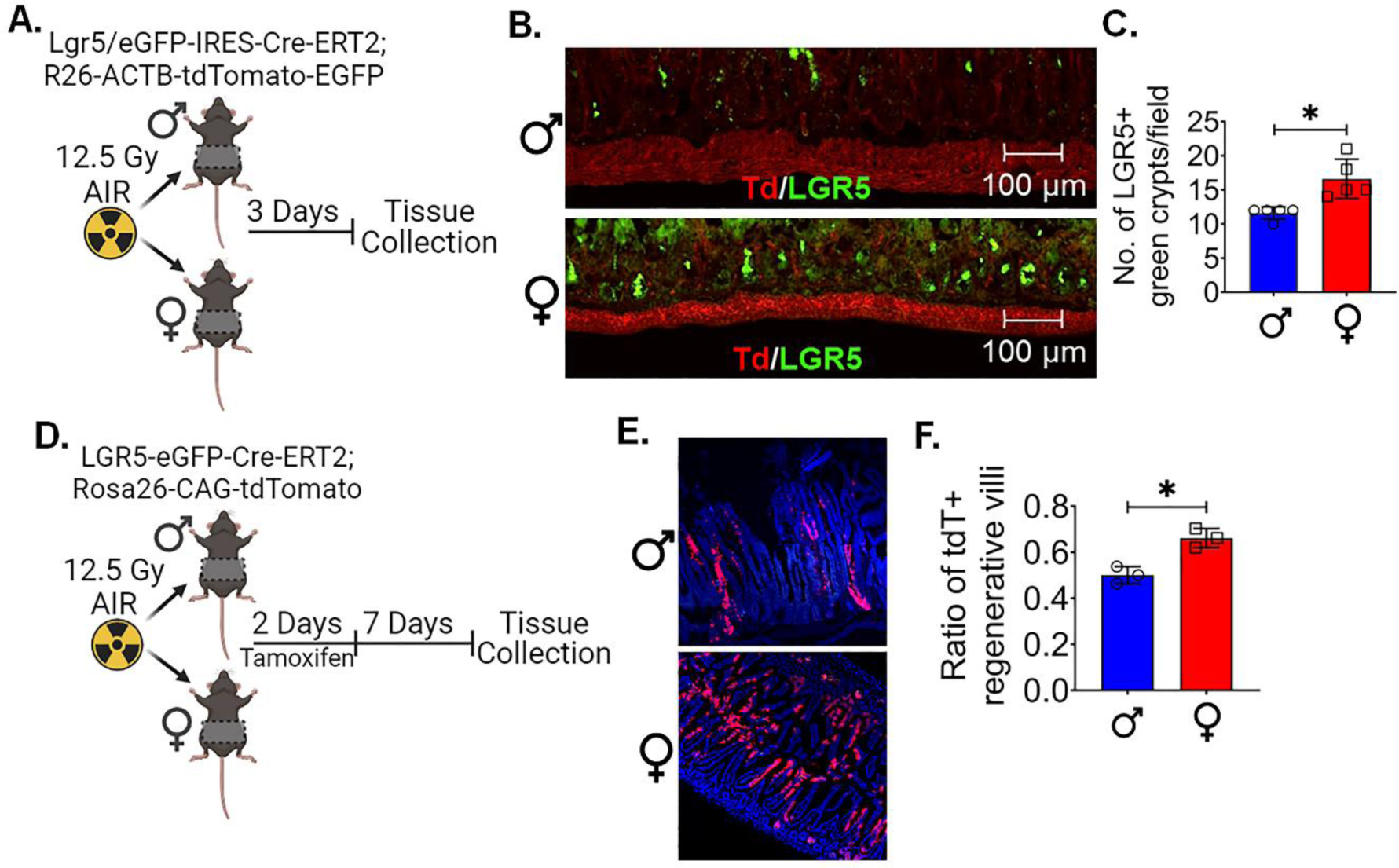
Female LGR5+ cells are radio-resistant and possess regenerative potential: A. Schematic representation of experimental design to analyze the survival of LGR5+ cells using Lgr5/eGFP-IRES-Cre-ERT2; R26-ACTB-tdTomato-EGFP mice following irradiation. B, C. Three days post-irradiation, a histological analysis shows a significant difference in the number of crypts containing LGR5+ cells in females compared to males. D. Schematic representation of experimental design to investigate the regenerative capacity of surviving ISC’s, a lineage tracing assay was performed using Lgr5-EGFP-ires-CreERT2-R26-CAG-tdT mice following irradiation. E, F. histological analysis shows, following irradiation and tamoxifen treatment, female intestinal epithelium revealed a significantly higher number of tdT-positive cells in regenerative villi compared to male mice. Data presented as the mean ± SD. (Significant level, *: p<0.05).

### Female LGR5+ cells are more capable of proliferation following radiation

After determining that female LGR5+ cells are more resistant to radiation, we wanted to assess if the surviving ISCs were capable of regeneration by performing *in vivo* lineage tracing assay using Lgr5-EGFP-ires-CreERT2-R26-CAG-tdT mice was performed. In this mouse tamoxifen-mediated activation of cre-recombinase under the Lgr5 promoter expresses tdTomato in epithelial cells derived from Lgr5-positive ISCs. Therefore, quantification of these tdTomato (tdT)-positive cells in irradiated epithelium in male and female mice determines the regenerative response of Lgr5-positive ISCs. Mice were exposed to 12.5Gy AIR, followed by Tamoxifen treatment demonstrated the presence of tdT-positive cells in the crypt epithelium in male and female mice intestinal tissue (Figure 2D). However, female intestinal epithelium shows a significantly higher number of tdT positive cells in regenerative villi compared to the male mice (p<0.05) (Figure 2E, F). All this evidence clearly demonstrates that female mice LGR5+ are more resistant to radiation toxicity and superior to promote intestinal epithelial regeneration by inducing the growth and proliferation of ISCs.

### Female intestinal epithelium shows less oxidative stress, mitochondrial biogenesis and better stemness

To investigate the underlying factors contributing to the observed variations in radiosensitivity between male and female mice intestinal epithelium, we examined oxidative stress gene expression analysis in male and female mice intestinal epithelial cells (IECs). 5 week and 12 weeks old male and female mice were exposed to 12.5Gy AIR and IECs were isolated after 4 hours of radiation exposure to analyze the reactive oxygen species (ROS) responsive gene (Figure 3A, C). The qPCR analysis of 5 week old male mice show significantly higher oxidative stress genes expression for Gclc (p<0.00005), Prdx-1 (p<0.00005), Nrf-2 (p<0.00005) and NqO1 (p<0.00005) in comparison to the female mice (Figure 3B). Interestingly, 12 week old male mice were also showing the similar pattern of oxidative stress genes expression for Gclc (p<0.005), Prdx-1 (p<0.05), Nrf-2 (p<0.005) and NqO1 (p<0.005) in comparison to the female mice (Figure 3D). As a significant amount of radiation induced DNA damage is transmitted by oxidative stress (38), these findings highlight a cause for the observed differences in radiation sensitivity. Interestingly, the western blot analysis of IECs also shows lower expression of some of the OXPHOS proteins (Figure 3E) in female mice compared to the male mice IECs. It is well documented that radiation upregulated the expression of genes related to OXPHOS (39) and mitochondrial biogenesis (40). Our quantitative PCR (qPCR) analysis of IECs from both male and female mice confirmed these findings, indicating that the expression of mitochondrial biogenesis genes, specifically NRF-1 and TFAM, was notably significantly lower in non-irradiated (p<0.05, p<0.05 respectively) (Figure 3F) and irradiated (p<0.005, p<0.00005 respectively) (Figure 3G) female mouse IECs. These results collectively suggest a reduced capacity for mitochondrial biogenesis in female mice.

**Figure 3:**
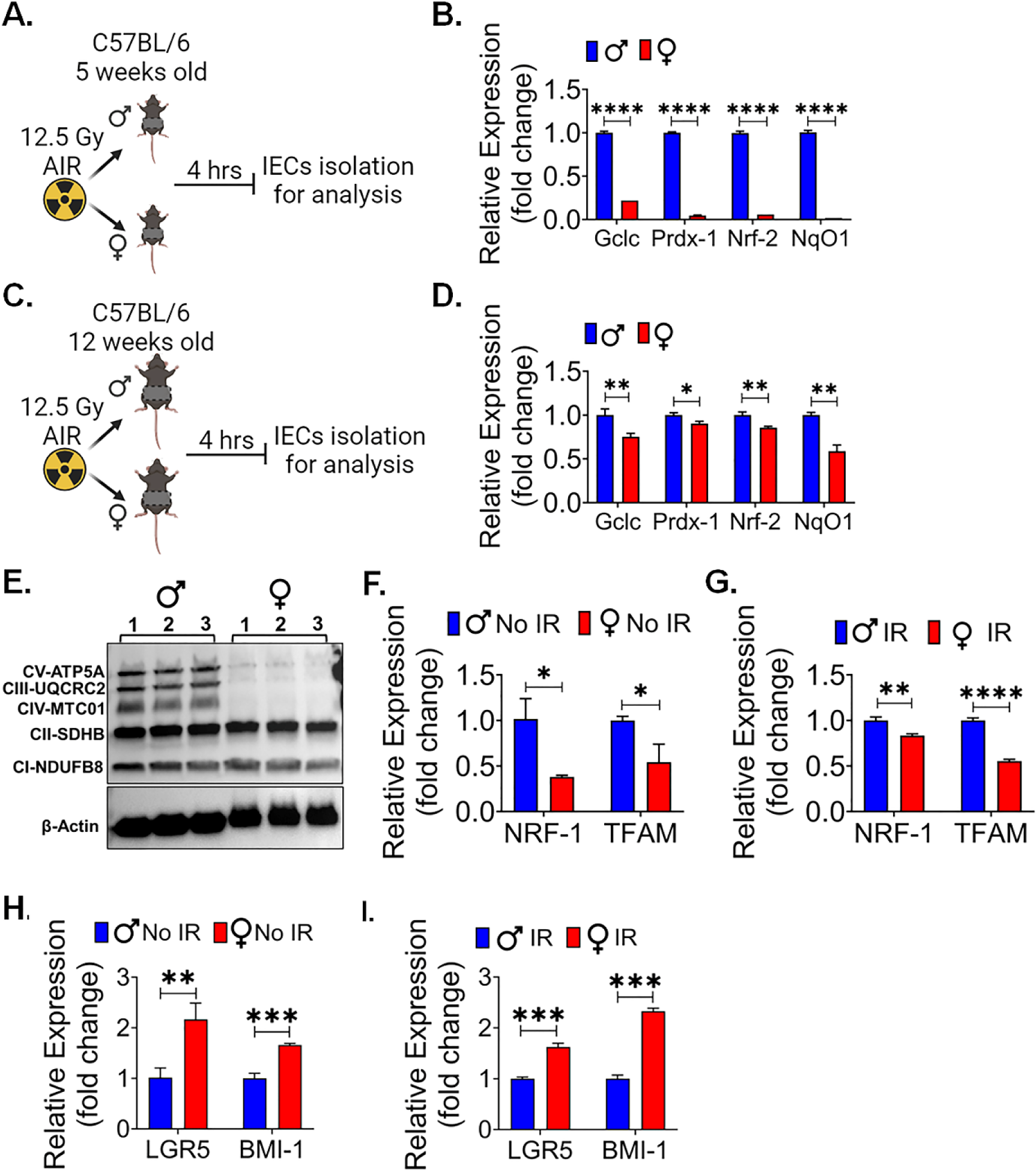
Female intestinal epithelium shows less oxidative stress, mitochondrial biogenesis and better stemness: Schematic representation of experimental design using A. 5-week-old and C. 12-week-old male and female mice, for analyzing the expression of oxidative stress genes following radiation exposure. B, D. comparative analysis of oxidative responsive genes expression (Gclc, Prdx-1, Nrf-2, and NqO1) analysis in 5-week and 12-week-old shows female have less oxidative stress responsive gene compared to male mice after radiation exposure. E. Irradiated female IECs shows less OXPHOS proteins expression, which plays a crucial role in mitochondrial function. F, G. qPCR analysis of mitochondrial biogenesis genes (NRF-1 and TFAM) in non-irradiated and irradiated IECs shows female have less mitochondrial biogenesis genes expression compared to male. H, I. qPCR analysis of major stem cell markers, LGR5 and BMI expression, in non-irradiated and irradiated IECs demonstrated female IECs have significantly higher expression of both the LGR5, and BMI-1 marker expression compared to male mice. Data presented as the mean ± SD. (Significant level, *: p<0.05, **: p<0.005, ***: p<0.0005, ****: p<0.00005).

Prior studies demonstrated the role of decreased OXPHOS and ROS production in enhancing the stemness in IECs (41, 42). Importantly, ISCs often rely on glycolysis as their primary energy source, which may contribute to the maintenance of their stem cell characteristics (43). The observed reduction in mitochondrial biogenesis in female IECs may potentially redirect cellular metabolism toward a more glycolytic profile, thereby preserving their stemness (44). Therefore, we also examined the major stem cells marker LGR5 and BMI expression in IECs, the qPCR analysis shows that of female IECs have significantly higher expression of LGR5 and BMI in non-irradiated (LGR5 (p<0.005) and BMI (p<0.0005) (Figure 3H) and irradiated (LGR5 (p<0.0005) and BMI (p<0.0005)) (Figure 3I) IECs cells compared to the male IECs. All these observations suggests that reduced levels of these oxidative events and mitochondrial biogenesis might support stemness of ISCs in females and, therefore, demonstrated resistance against radiation than males.

### Female intestinal epithelium shows less radiation induced DNA damage

Radiation exposure is known to induce DNA damage in various tissues, including the sensitive intestinal epithelium. Gamma-H2AX (γ-H2AX) is a phosphorylated form of the histone protein H2AX and is known to form distinct foci at sites of DNA double-strand breaks (DSBs), which are a prevalent and critical form of DNA damage triggered by ionizing radiation (45). To assess and compare the response to radiation-induced DNA damage between male and female mice, particularly following exposure to a significant dose of 12.5 Gy of PBI, immunofluorescent staining for γ-H2AX was performed in the intestinal tissue sections to visualize and quantify the extent of DNA damage in the intestinal epithelium of both male and female mice. This approach allows for a precise assessment of potential sex-based differences in the response to radiation-induced DNA damage within the intestinal tissue. The immunofluorescent staining shows no significant differences for γ-H2AX staining in between non-irradiated male and female mice intestinal tissue (Figure 4A). However, a significantly higher number of γ-H2AX foci was observed in irradiated male mice in comparison to their female counterparts (Figure 4A, B). This finding indicates that, following irradiation, male mice exhibited a greater extent of radiation-induced DNA damage, specifically in the form of γ-H2AX foci formation. These difference in DNA damage may contribute to the observed differences in radio-sensitivity between male and female mice.

**Figure 4:**
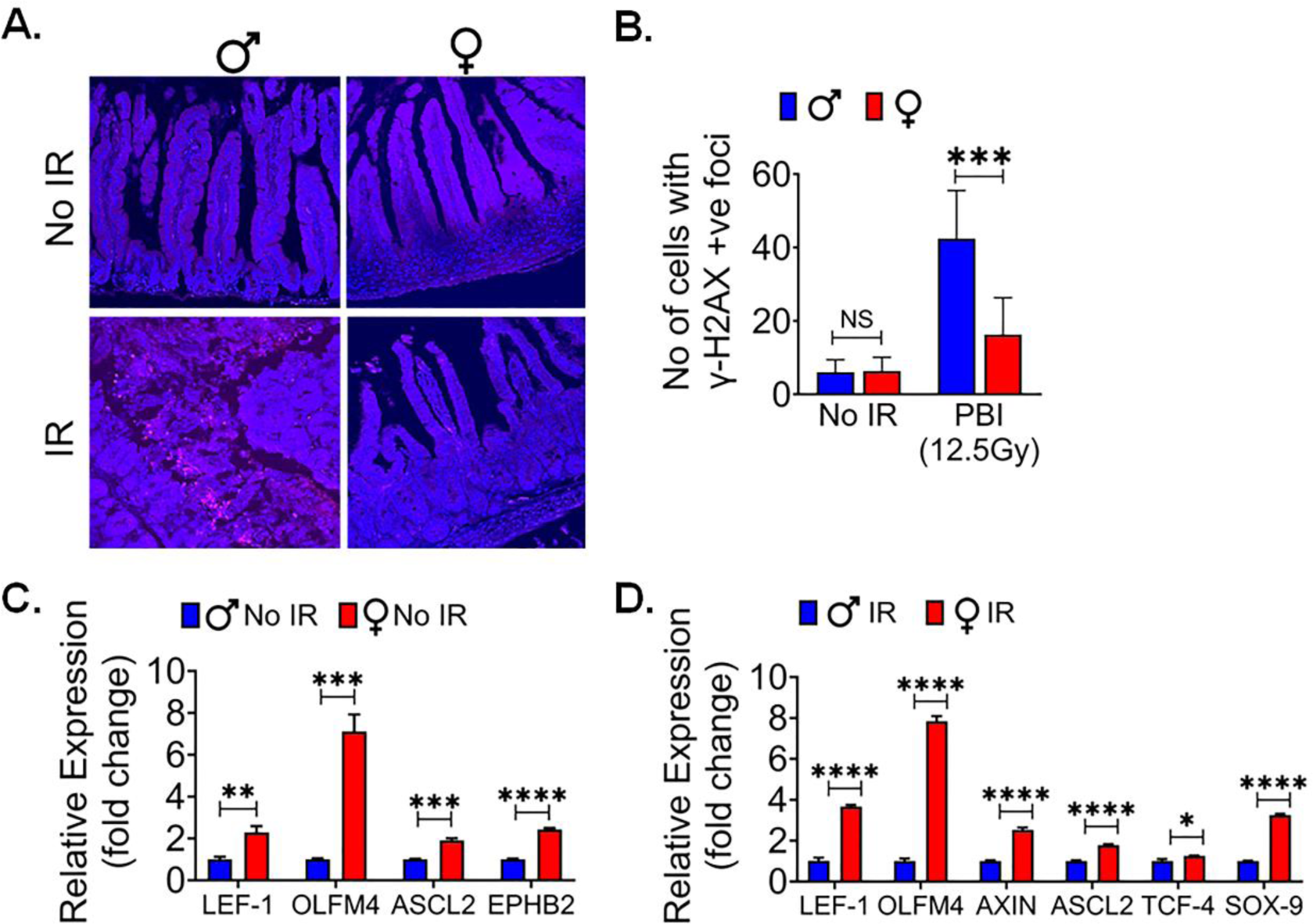
Female intestinal epithelium is resistant to radiation induced DNA damage. A. shows immunofluorescent staining for γ-H2AX in non-irradiated and irradiated male and female mice intestinal tissue. B. shows no significant differences in γ-H2AX foci between non-irradiated male and female mice, while irradiated male mice intestinal tissue shows significantly higher number of γ-H2AX foci compared to their female counterparts. C, D. The qPCR analysis of β-catenin target genes in non-irradiated and irradiated IECs from both female and male mice, revealed a significant upregulation of multiple β-catenin target genes in female IECs compared to male IECs in both non-irradiated and irradiated mice IECs. Data presented as the mean ± SD. (Significant level, *: p<0.05, **: p<0.005, ***: p<0.0005, ****: p<0.00005, NS: not significant).

Canonical WNT signaling is key for intestinal epithelial repair and regeneration. We reported earlier that pharmacological activation of WNT β-catenin signaling can promote intestinal epithelial repair and regeneration and mitigate radiation-induced intestinal injury (31, 46). To understand the role of WNT β-catenin signaling, we conducted qPCR analysis of WNT target genes in IECs from female and male mice. We found a significant upregulation of multiple β-catenin target genes in female IECs when compared to male IECs, both in non-irradiated (Figure 4C) and irradiated (Figure 4D) conditions. These findings suggest that the activation of WNT β-catenin signaling is more distinct in female IECs, highlighting a potential sex-specific difference in the response to WNT pathway activation in the context of intestinal epithelial radiosensitivity.

### Female ex-vivo intestinal organoid is more radio-resistant

Organoids have become an established ex-vivo model of intestinal stem cells (47). To identify the local differences in LGR5+ cells between male and female mice ex-vivo intestinal organoid were cultured and subjected to various radiation doses and examined for their morphological features (Figure 5A). Specifically, we focused on assessing the organoids’ ability to regenerate, as indicated by the budding phenomenon. Our observations revealed a significant difference in the budding crypt ratio between male and female intestinal organoids when exposed to radiation doses ranging from 0 to 8Gy (Figure 5B). Furthermore, we observed a significant increase in the average cross-sectional area of female organoids, particularly at higher radiation doses of 6 and 8 Gy, in comparison to their male counterparts. However, no significant differences were detected at radiation doses below 6Gy (Figure 5C, D). These data show the morphological disparities within organoids derived from male and female mice exposed to varying radiation doses yielded an intriguing finding. A notable difference between the two groups became evident at a radiation dose of 6Gy. Therefore, further experiments focus exclusively on the 6Gy radiation dose.

**Figure 5:**
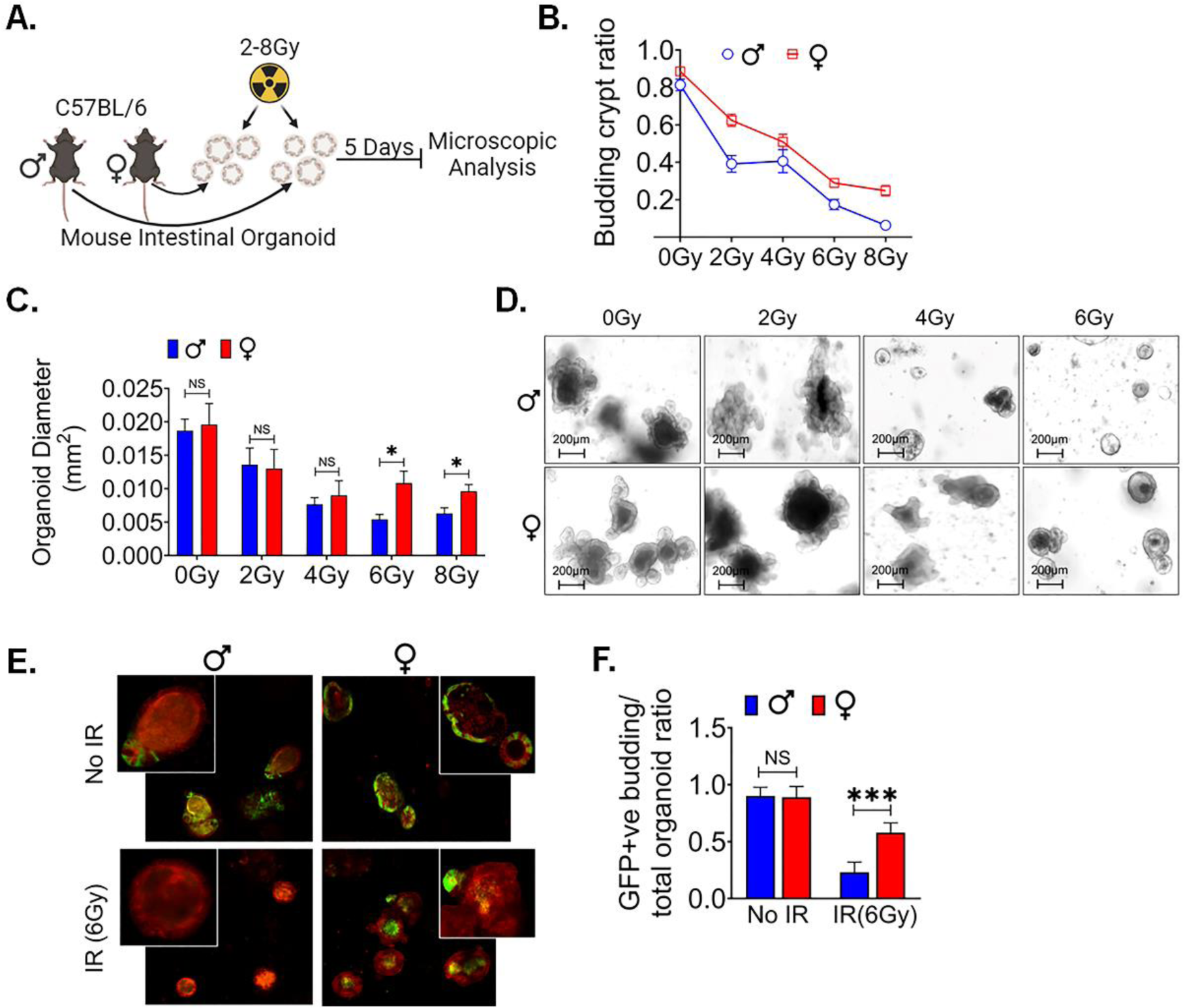
Female ex-vivo intestinal organoid culture is more radio-resistant. A. Schematic representation of male and female mice intestinal organoids culture, radiation exposure for examining their morphological features. B. shows higher budding crypt ratio in female intestinal organoid compared to male following exposure to 0 to 8 Gy of irradiation. C, D. showing assessment of the average cross-sectional area of female organoids, with noticeable increases, especially at higher radiation doses (6 and 8 Gy), when compared to male organoids. No significant differences were detected at radiation doses below 6 Gy. E. shows organoids derived from Lgr5/GFP-IRES-Cre-ERT2; R26- ACTB-tdTomato-EGFP knock-in mice. F. shows no significant differences in the ratio of GFP+ budding organoids to total organoids ration in non-irradiated male and female organoids. However, within 48 hours post-irradiation, female mice organoids exhibited a significantly higher GFP+ budding organoid/total organoid ratio compared to male mice organoids. Data presented as the mean ± SD. (Significant level, *: p<0.05, ***: p<0.0005, NS: not significant).

We repeated this study in organoids derived from Lgr5/GFP-IRES-Cre-ERT2; R26- ACTB-tdTomato-EGFPknock-in mice where LGR5+ ISCs can be detected based on EGFP. In non-irradiated conditions, we observed no significant difference in the ratio of GFP+ budding organoids to total organoids between male and female organoids. However, a noticeable sex-based difference was emerged within 48 hours post-irradiation in male organoids. We notice a reduction in the population of Lgr5+ve ISCs, resulting in a significant loss of GFP+ budding crypts with alterations in the morphology of existing crypts, indicative of the inhibition of ISC growth and differentiation in response to the 6 Gy radiation exposure. In contrast female mice organoids exhibited a significantly higher GFP+ budding organoid/total organoid ratio, highlighting their radio-resistance property (p<0.0005) (Figure 5E, F). These findings highlight that female mice ISCs are more resistant radiation-induced injury.

### Female intestinal organoid mimics the in-vivo characteristics

Further we examined the comparison between male and female organoids in DNA damage markers following irradiation (Figure 6A). We observed that DNA damage marker in mouse intestinal organoid exposed to irradiation shows a significant difference between male and female organoid. At a radiation dose of 6 Gy, we observed a significantly higher number of γH2AX foci in male mice intestinal organoids compared to female mice (p<0.005) (Figure 6B, C).

**Figure 6:**
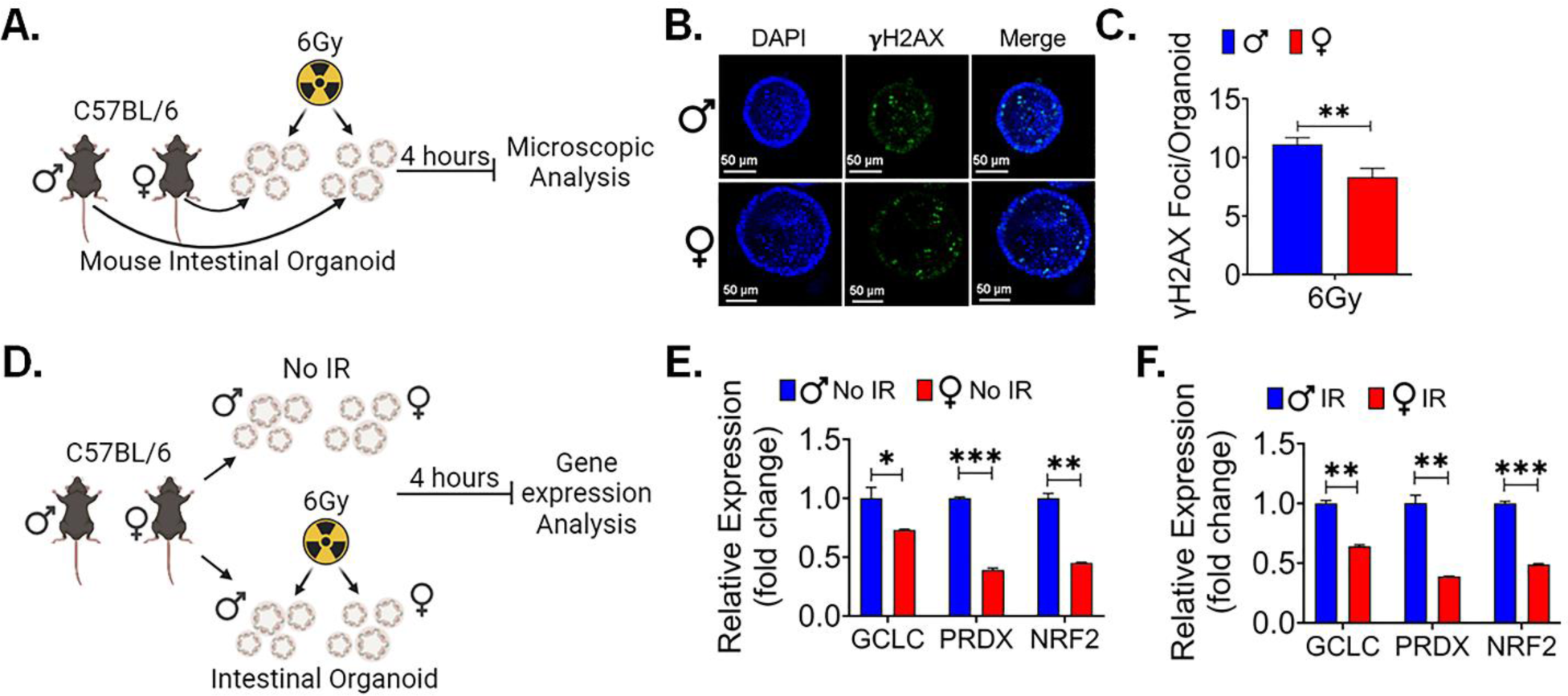
Female intestinal organoids are resistant to radiation induced DNA damage: A. Schematic representation of male and female mice intestinal organoids culture, radiation exposure for microscopic analysis. B, C. shows significantly higher number of γH2AX foci in male mice intestinal organoids when compared to female mice, highlighting sex-based differences in the response to DNA damage induced by radiation exposure. D. Schematic representation of male and female mice intestinal organoids culture, radiation exposure for gene expression analysis. E, F. The qPCR analysis shows significantly upregulated genes of oxidative stress responsive gene Gclc, Prdx, and Nrf2 expression in non-irradiated and irradiated male organoids when compared to female mice organoids. Data presented as the mean ± SD. (Significant level, *: p<0.05, **: p<0.005, ***: p<0.0005).

Additionally, when examining oxidative stress gene expression, consistent with our in-vivo findings, we found a significant difference in expression profiles. qPCR analysis of oxidative stress markers shows significantly upregulated gene expression of Gclc (p<0.05, p<0.005 respectively), Prdx (p<0.0005, p<0.005 respectively) and Nrf2 (p<0.005, p<0.0005 respectively) in non-irradiated and irradiated male organoids in comparison to the female mice organoids (Figure 6D-F).

All these ex-vivo findings collectively suggest that differences in the ability of LGR5+ cells to respond to radiation are not solely attributed to hormonal variations or distal organ interactions encountered in an in-vivo setting. Moreover, they pinpoint the male-specific surge in oxidative stress as a post-irradiation phenomenon, as male organoids displayed higher basal levels of oxidative stress even in the absence of radiation exposure.

### Human Organoids also show similar female Radio resistance

In order to validate the sex-specific differences observed in murine models, we sought to investigate whether these findings could be replicated in human tissue. To accomplish this, we utilized male and female surgical specimens obtained from normal colon regions, positioned at least 10 cm away from malignant sites (48). From these specimens, we generated ex-vivo crypt organoids and subjected them to radiation exposure at a dose level of 6 Gy (Figure 7A). Interestingly, in human male organoids, all budding crypts were found to have disappeared following exposure to 6Gy, in contrast to the human female organoids (Figure 7B). Furthermore, when assessing the radiation response of patient-derived organoids at various time points after exposure to 6 Gy, we found a remarkable observation. Female organoids exhibited a significantly larger cross-sectional area at both day 1 (p<0.05) and day 7 (p<0.005) post-exposure (Figure 7C). These results are very similar to what we observed in female murine organoids, thereby indicating a remarkable consistency in outcomes across species.

**Figure 7:**
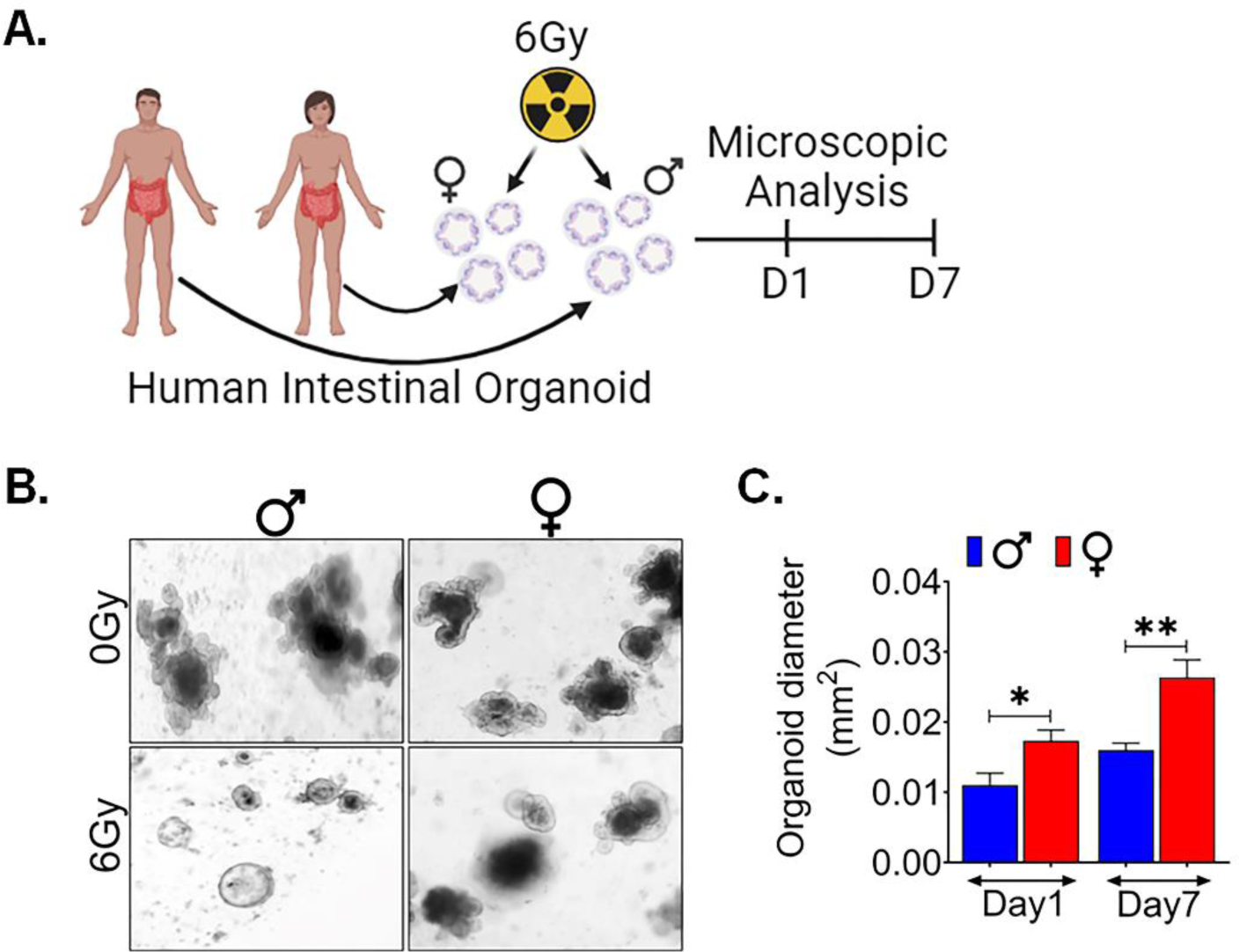
Human Organoids shows females are radio resistance: A. Schematic representation of human male and female intestinal crypt organoids culture, radiation exposure for their morphological features by microscopic examining. B, C. showing assessment of the average cross-sectional area of female organoids, was significantly higher at day 1 and 7 following radiation exposure, when compared to male organoids, highlighting a sex-based differences in the post-radiation responses. Data presented as the mean ± SD. (Significant level, *: p<0.05, **: p<0.005).

### RNAseq analysis of Human LGR5+ Cells Shows no DDR Male Female Difference, Suggests mitochondrial reason

To gain deeper understanding into the underlying factors contributing to disparities in radiosensitivity of the LGR5+ cell population between males and females, we conducted a comprehensive mRNA expression analysis using patient-derived organoids (PDOs). Specifically, PDOs were generated from tissue samples obtained from 10 patients, (5 males and 5 females) (Figure 8A). Once the organoid cultures were successfully established, they were dissociated into single-cell suspensions, and LGR5+ cells were isolated employing magnetic microbeads method. Subsequently, the isolated cells were processed for RNA sequencing analysis (Figure 8A).

**Figure 8:**
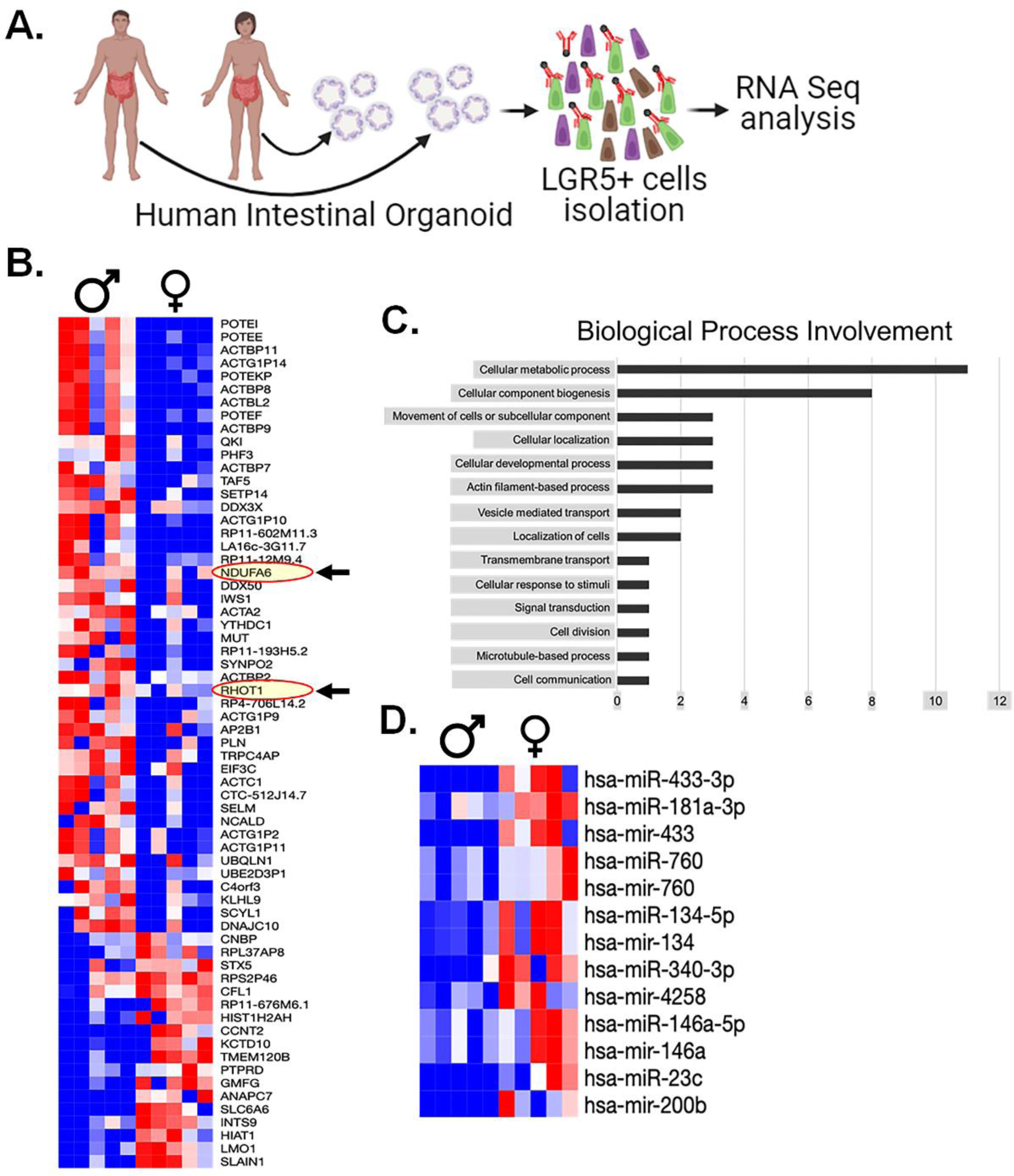
Transcriptomics analysis: A. Experimental Design for Patient-derived organoids culture, isolation of LGR5+ cells for RNA seq analysis. B. shows differential mRNA expression in male and female LGR5+ cells. RNA seq. analysis revealed significant differences in the expression of 55 mRNA genes, highlighting two mitochondrial proteins, NDUFA6 (an Electron Transport Chain protein) and RHOT1 (a mitochondrial membrane protein), were upregulated in males compared to females (shown in red oval with arrow), C. Gene Ontology Analysis of Differentially Expressed Genes to categorize the biological processes associated with the 55 differentially expressed genes, D. miRNA analysis of serum samples shows 13 miRNAs that exhibited statistically significant differences between males and females.

The RNAseq analysis revealed substantial disparities in the expression levels of 55 mRNA genes (Figure 8B). Interestingly, among these findings, we observed the upregulation of two mitochondrial proteins, namely NDUFA6 (an Electron Transport Chain protein) and RHOT1 (a mitochondrial membrane protein) in males compared to females (Figure 8B). To systematically categorize the implications of these mRNA findings, we subjected this gene list to a gene ontology analysis, resulting in the involvement of specific biological processes (Figure 8C).

Remarkably, one of the most notable findings we observed from this analysis is the absence of measurable differences in DNA damage response (DDR) machinery expression between the sexes. Instead, the predominant category pertains to metabolic processes, supporting with our previous observations of oxidative stress. This collective evidence points toward mitochondrial factors as the key determinants underlying our findings between male and female.

### miRNA analysis of human serum highlights several biomarkers

Finally, in an attempt to further explain any observed differences in human LGR5+ populations, we performed a miRNA analysis of serum samples from same male and female patients whose intestinal surgical samples used for the intestinal organoids. The miRNA analysis yielded a collection of 13 miRNAs that exhibited statistically significant distinctions across all males and females studied (Figure 8D). Interestingly, among these, several miRNAs had previously been explored for their potential roles in modulating radiosensitivity. Out of 2079 identified mi-RNA from the plasma of male and female, 37 mi-RNAs are associated with mitochondrial biogenesis (Supplementary Table 2 and 3). These finding shows that some of these miRNAs may play a role in modulating the mitochondrial biogenesis process and enhancing the response to radiation resistance, especially in females. Understanding the specific roles of these miRNAs is essential for understanding the mechanisms behind the observed sex-based differences in radio-resistance.

Nonetheless, hsa-mir-760, which displayed significantly higher expression levels in female serum samples (p<0.05). It is noteworthy that previous investigations have linked low hsa-mir-760 expression to increased cell proliferation in colorectal tissue in in-vitro (49). This interesting finding suggests that hsa-mir-760 may play a pivotal role in modulating cell proliferation, with potential implications for the observed differences in radiosensitivity.

Given these interesting findings, it is evident that further research is needed to investigate the individual functions of these miRNAs in the context of mitochondrial biogenesis and their impact on radioresistance. These studies will be crucial in unveiling the intricate molecular mechanisms that underlie the differences in radiosensitivity among human LGR5+ populations. The complexity of miRNA regulation and its multifaceted roles necessitate a deeper exploration to decipher the precise functions and contributions of these specific miRNAs in the context of radiosensitivity.

## Discussion

In this study, we investigate the potential disparities in the response of male and female intestinal epithelium to radiation exposure, as well as to unravel the underlying mechanisms contributing to these differences. The results of our experiments demonstrate that female intestinal tissue exhibits a remarkable resistance to radiation-induced toxicity, at both histological and molecular levels. Notably, we identified a significant radiosensitivity difference in male and female intestinal epithelium and LGR5+ stem cell population, which was consistently identified in both murine and human subjects. Specifically, female LGR5+ stem cells displayed enhanced radioresistance and capable of proliferation following radiation exposure. This robustness of female LGR5+ ISCs is of particular importance since ISCs play a pivotal role in preserving the homeostasis and regenerative potential of intestinal tissue. Interestingly, our study suggested that the distinct response to radiation is not due to DNA damage response machinery but rather to metabolic and mitochondrial distinctions between the sexes.

Our molecular analysis has uncovered an array of factors that collectively account for the sex-based differences in radiosensitivity observed in the intestinal epithelium. These findings provide an understanding of the molecular differences underlying the remarkable radio resistance exhibited by female. Notably, these factors involve oxidative stress, mitochondrial biogenesis, cellular metabolism, and stemness markers. It is well reported that radiation response upregulates the oxidative stress (50, 51). Which is one of the key factors we also observed in our study. Our analysis of oxidative stress gene expression shows increased oxidative stress in male intestinal epithelial cells following irradiation compared to female. This upregulation in oxidative stress, especially after radiation exposure, is a critical factor contributing to the enhanced sensitivity of male mice to radiation-induced damage than female mice. In contrast, female mouse intestinal epithelial cells display lower expression of oxidative phosphorylation (OXPHOS) proteins and genes associated with mitochondrial biogenesis, specifically NRF-1 and TFAM. This observation suggests that female intestinal epithelial cells undergo a metabolic shift away from OXPHOS and towards glycolysis. This metabolic modulation supports with the concept that glycolysis supports stemness in cells (52). This shift potentially enhances the maintenance of stem cell characteristics (53) in female subjects, providing an additional layer of protection against radiation-induced injury. This observation is supported by the notably higher expression of major stem cell markers, including LGR5 and BMI1, in female intestinal epithelial cells. These findings collectively emphasize the stemness maintained by female cells, which contributes to their radioresistance properties.

It is worth highlighting that the modulation of metabolic pathways, including the shift towards glycolysis, has been recognized as a strategy employed by various stem cell populations to sustain their undifferentiated state (54, 55). In our study considering this concept to LGR5+ population of intestinal epithelial cells to maintain their stemness and regenerative capacity in the presence of radiation-induced stress is a key factor contributing to the overall radio resistance of the female intestinal epithelium. The immunofluorescent staining for γ-H2AX indicated that male mice exhibited a greater extent of radiation-induced DNA damage. These findings contribute to the overall understanding of why male mice are more susceptible to radiation-induced injury. Moreover, the activation of WNT β-catenin signaling was more significant in female intestinal epithelial cells, indicating a potential sex-specific difference in the response to WNT pathway activation, which is vital for tissue repair and regeneration.

The findings of our study demonstrate convincing evidence for the radioresistance of ex-vivo intestinal organoid cultures from female mice. These organoids have emerged as a valuable model for investigating the biology of intestinal stem cells, and our experiments provided critical insights into sex-based differences in radiation response. Specifically, our data indicate that female organoids exhibit remarkable radioresistance, with a significantly higher budding crypt ratio, especially at higher radiation doses of 6 and 8Gy. In contrast, male organoids displayed pronounced sensitivity to radiation, as evidenced by the reduction in budding crypts and alterations in crypt morphology following exposure to 6Gy. These sex-based distinctions became notably with more than 6Gy radiation dose. Moreover, the use of genetically modified mouse Lgr5/GFP-IRES-Cre-ERT2; R26-ACTB- tdTomato-EGFPknock-in allowed us to visualize and quantify LGR5+ intestinal stem cells based on EGFP expression which shows a sex-specific differences within 48 hours post-irradiation. These differences can become crucial for considering radiation therapy and offer potential strategies for enhancing the protection of healthy tissues during such treatments.

Our findings using human organoids supported the observation made in murine models, highlighting the consistency of female radio resistance. These human organoid models also provided insights into the sex-specific differences in DNA damage markers following irradiation. The RNA sequencing analysis of human LGR5+ cells revealed differences in the expression of several genes, notably the upregulation of mitochondrial proteins in males, suggesting the involvement of mitochondrial factors as key determinants of radiosensitivity. Importantly, this analysis did not show measurable differences in DNA damage response machinery between the sexes. The miRNA analysis of human serum samples identified several miRNAs with statistically significant distinctions between males and females. The identified miRNA may play a critical role in modulating cell proliferation and could contribute to the observed differences in radiosensitivity.

In summary, the findings of this study provide a comprehensive understanding of the sex-based differences in radiosensitivity to intestinal epithelium. These findings collectively show a remarkable radioresistance of female intestinal tissue. These findings have important implications for radiation therapy and highlight the need for personalized treatment strategies based on sex-specific differences in radiosensitivity. Further research into the specific roles of the identified genes, proteins, and miRNAs will be essential for a deeper understanding of the molecular mechanisms driving these differences and their potential clinical applications.

## Author Contributions

Conceptualization S.S.; methodology, S.S.; P.B.; RZ; R.M.C; formal analysis, R.Z., R.M.C. P.B.; investigation, R.Z., R.M.C. P.B.; resources, S.S.; data curation, R.Z., R.M.C. P.B.; writing— R.Z., R.M.C, S.S.; review and editing, S.S. R.Z., R.M.C.; supervision, S.S.; funding acquisition, S.S. All authors have read and agreed to the published version of the manuscript.

## Funding

This work was supported by grants from NIH U01AI170036 (S.S.).

## Institutional Review Board Statement

All animal studies were performed under the guidelines and protocols of the Institutional Animal Care and Use Committee of the University of Kansas Medical Center. All the animal experimental protocols were approved by the Institutional Animal Care and Use Committee of the University of Kansas Medical Center (ACUP number 2019-2487).

## Data Availability Statement

Data is contained within the article and Supplementary Material.

## Acknowledgments

The authors are grateful to Carlton Anderson and Gabrielle Cannon from UNC Advanced Analytics (AA) Core facility for providing a material method section for RNA seq analysis performed at their Advanced Analytics (AA) Core facility.

## Conflicts of Interest

The authors declare no conflict of interest.

---

**Supplementary Table-1.**
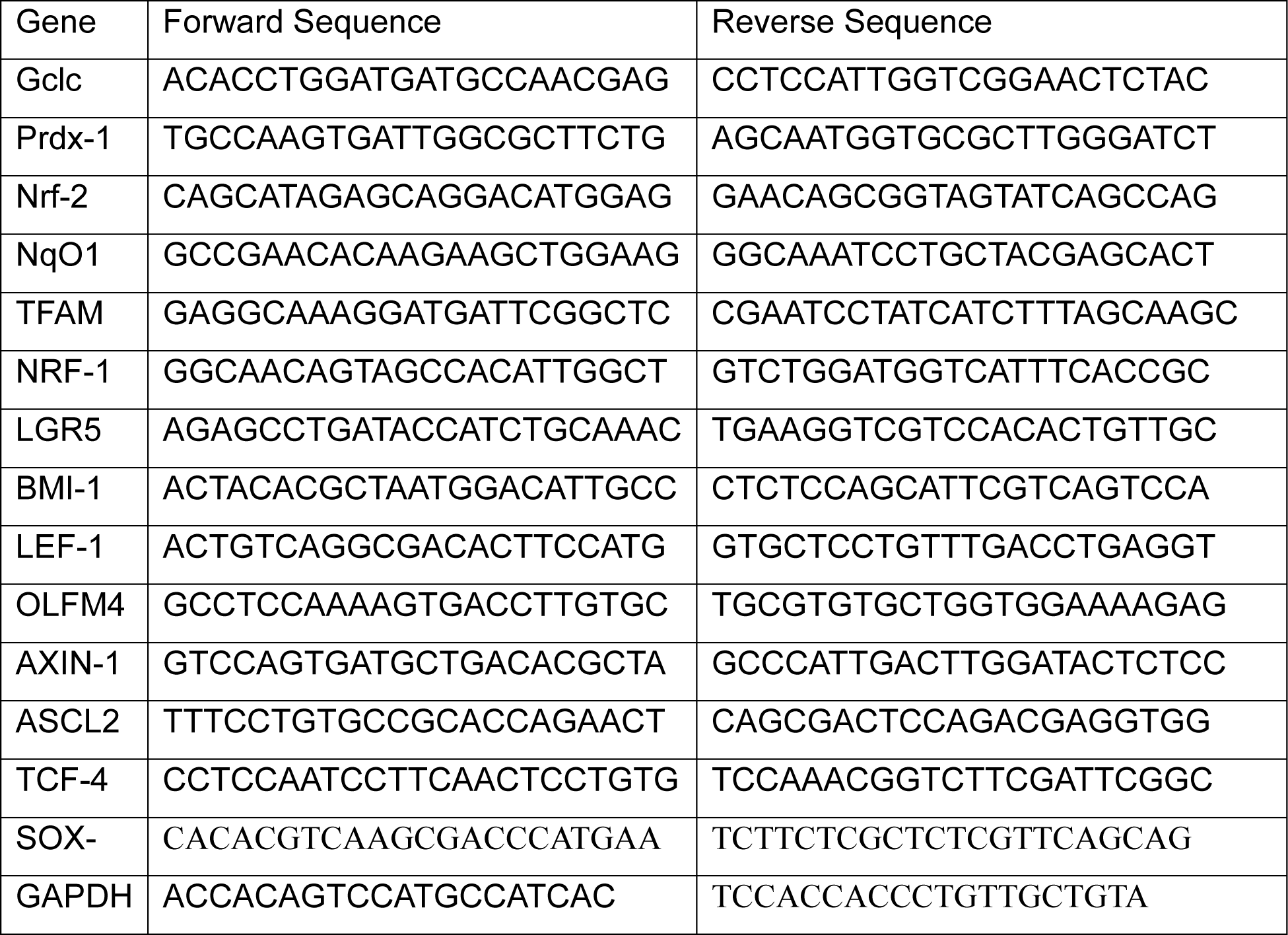
Primer Sequences.

**Supplementary Table-2.**
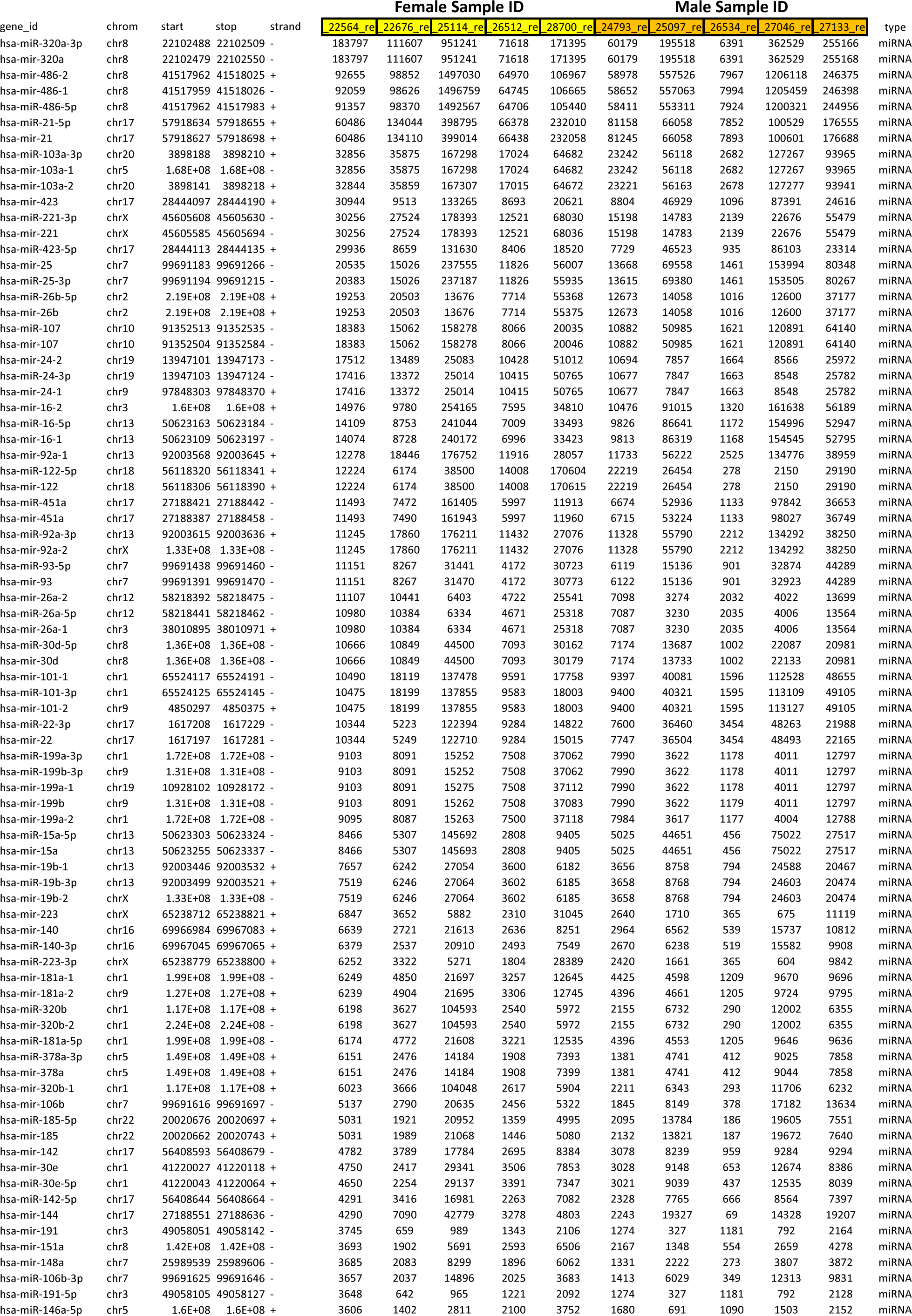
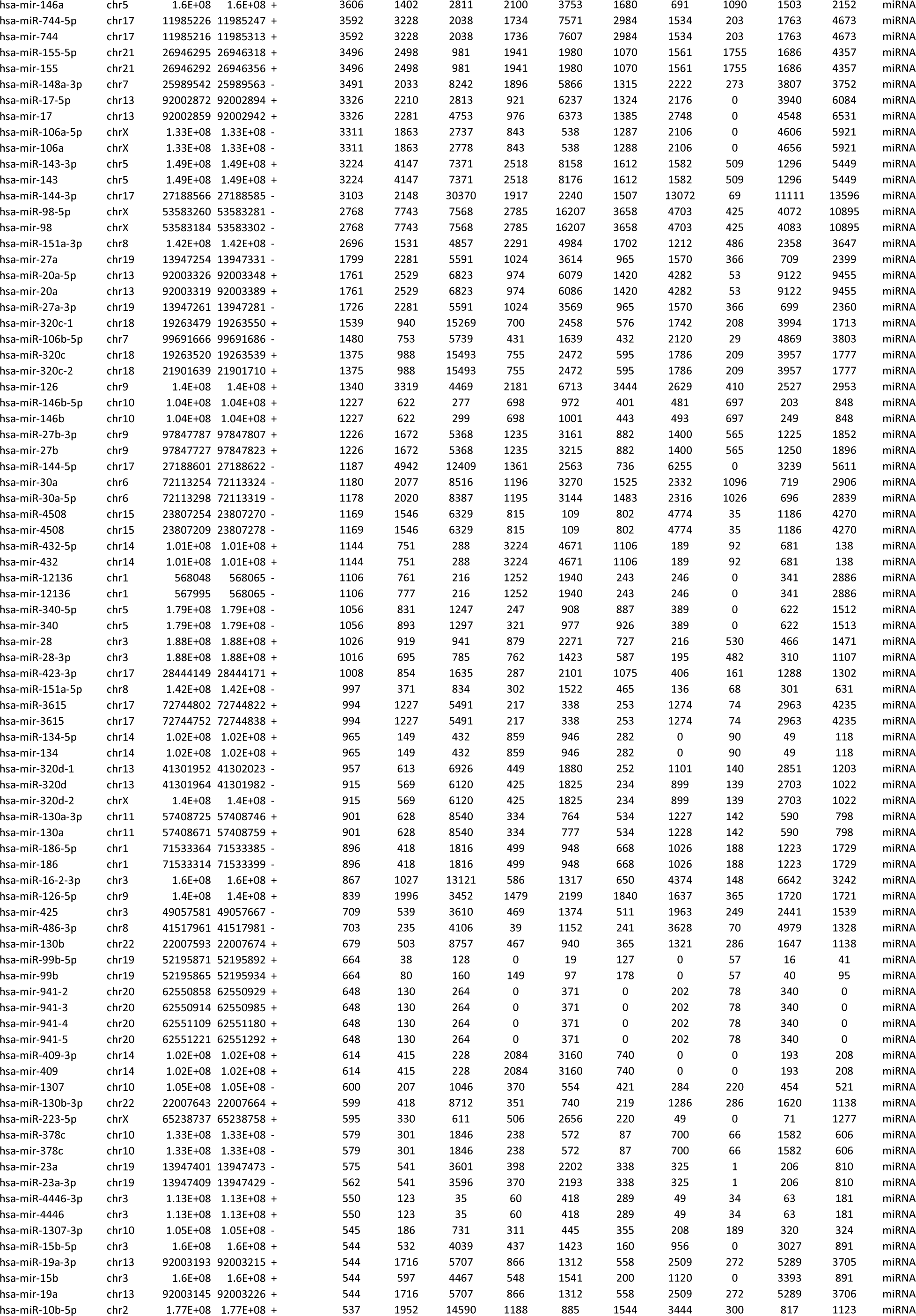
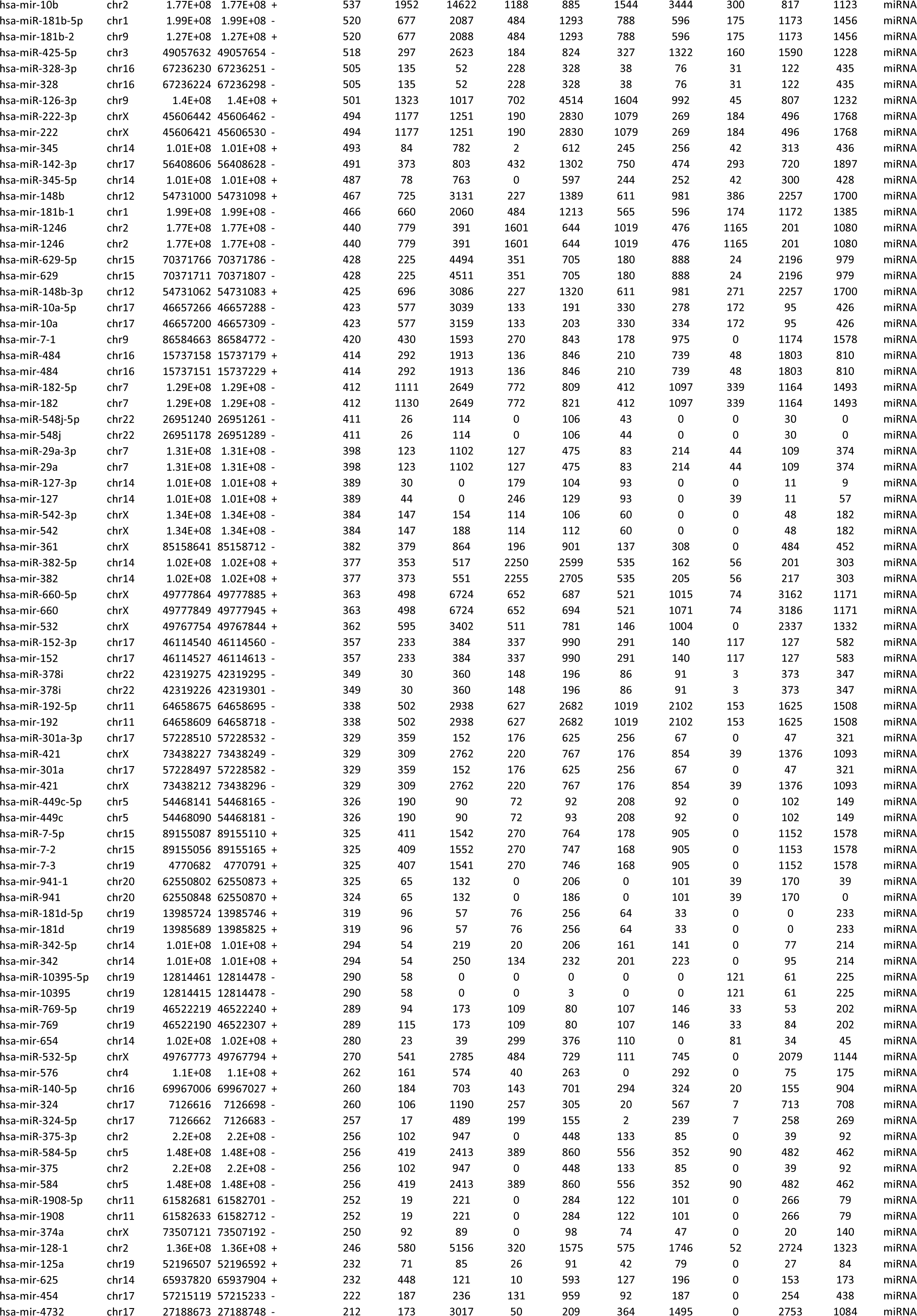
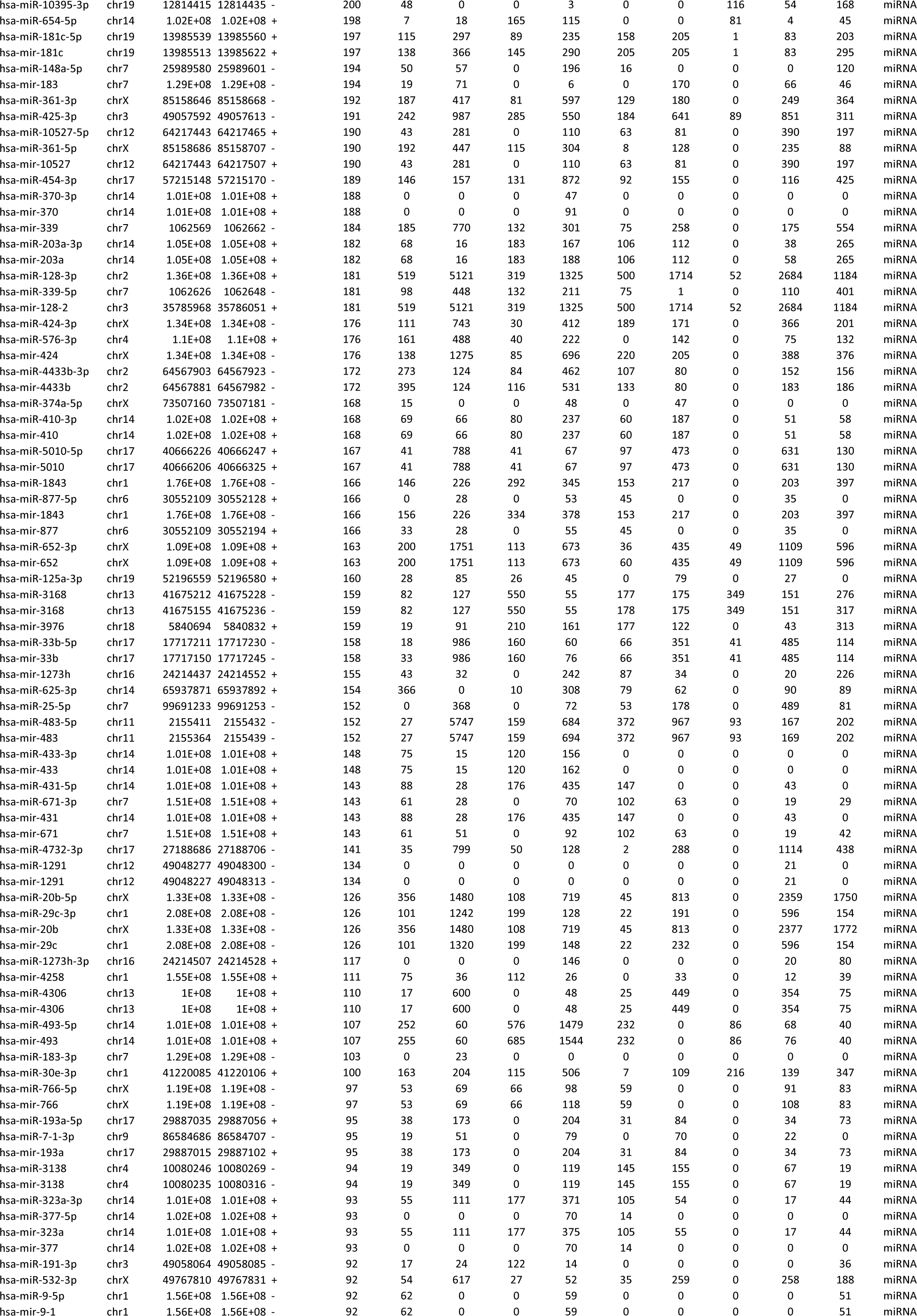
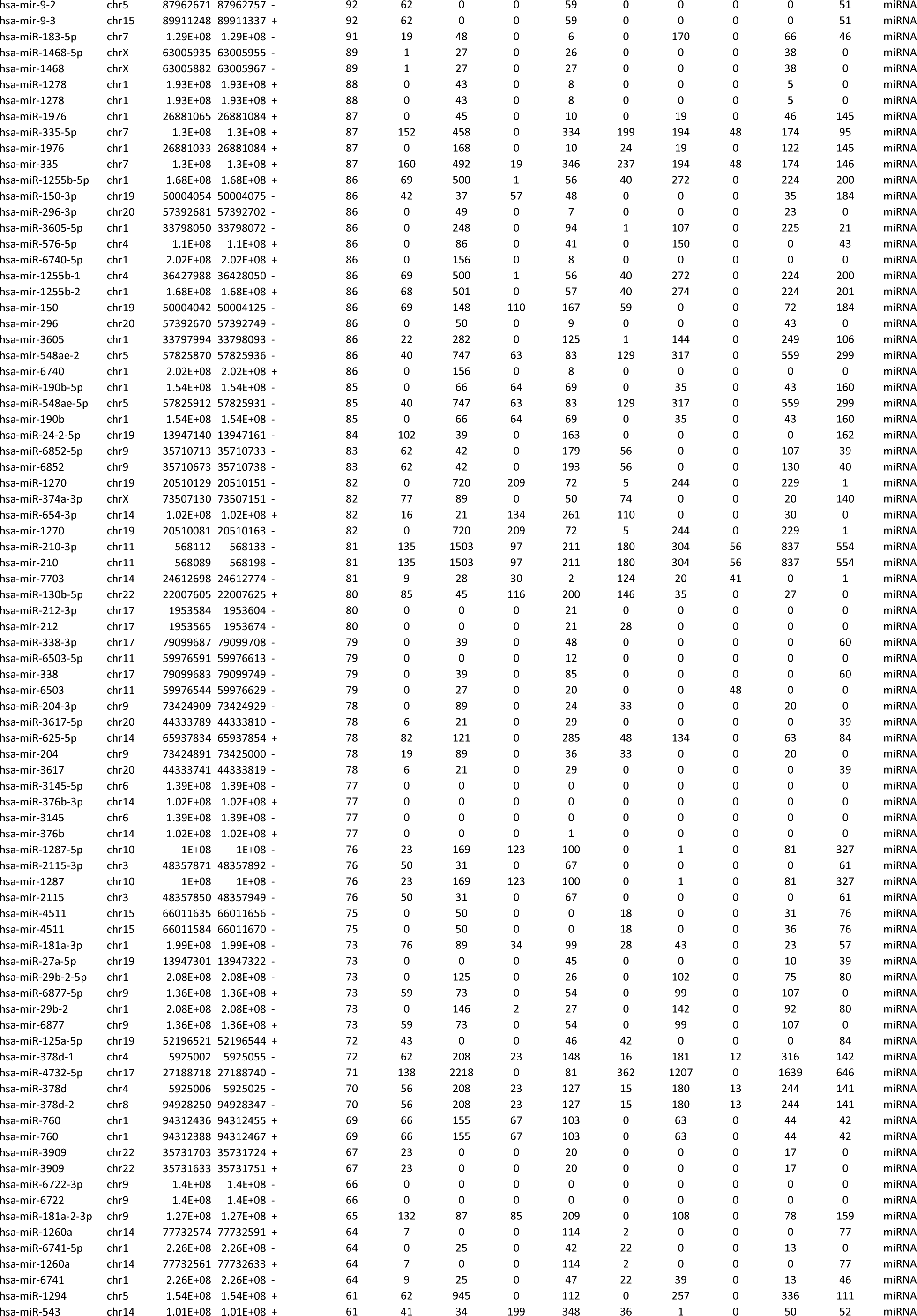
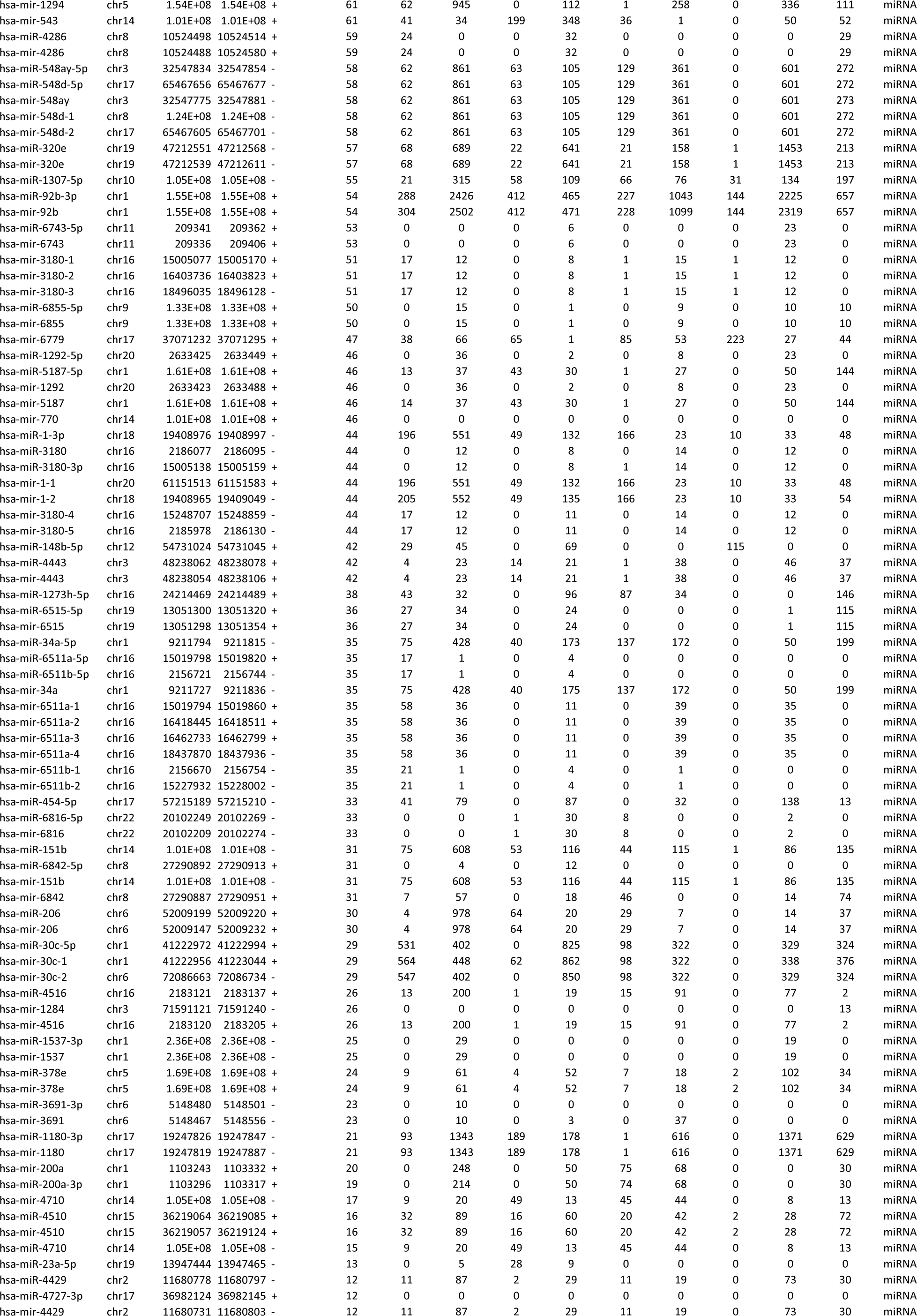
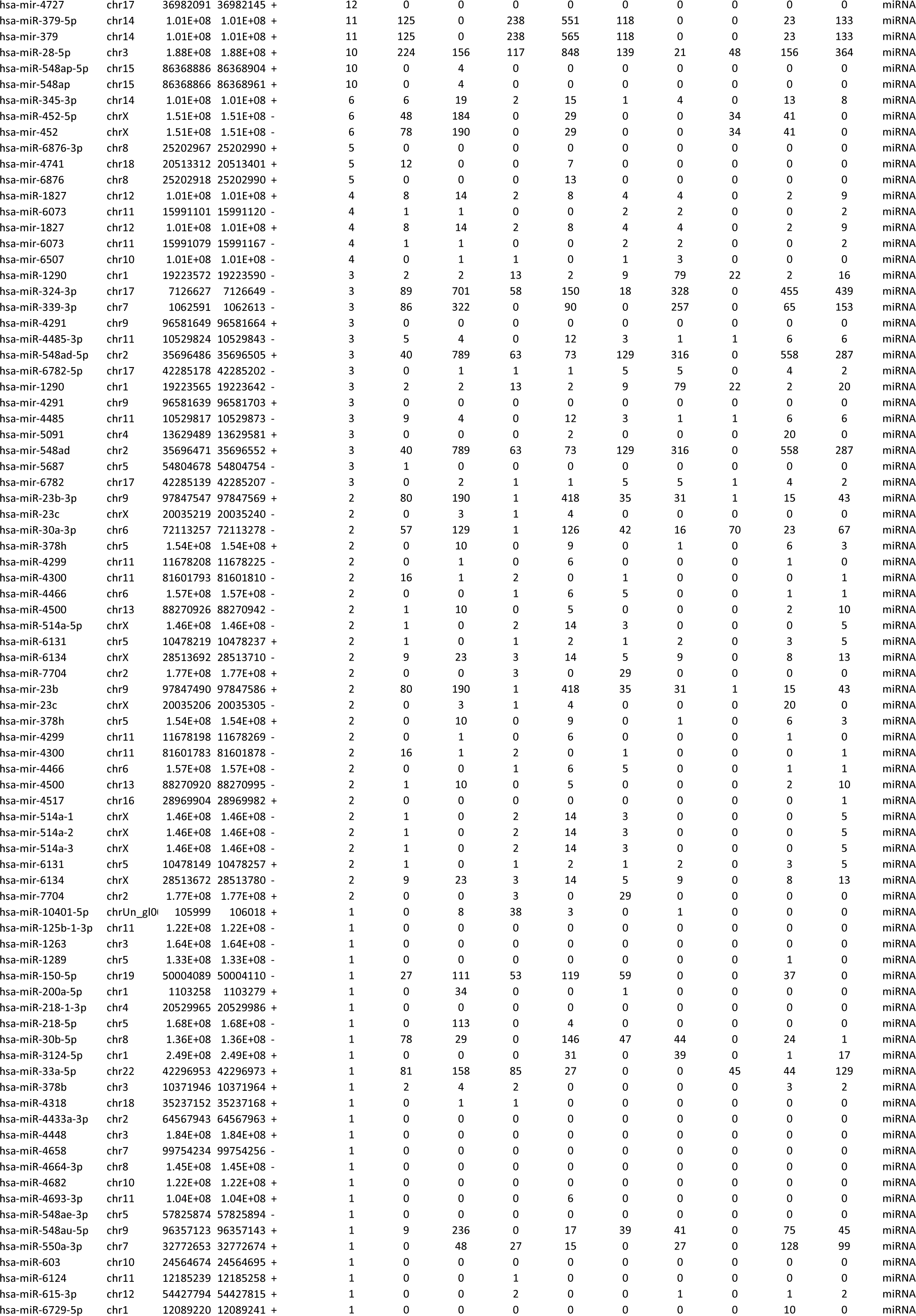
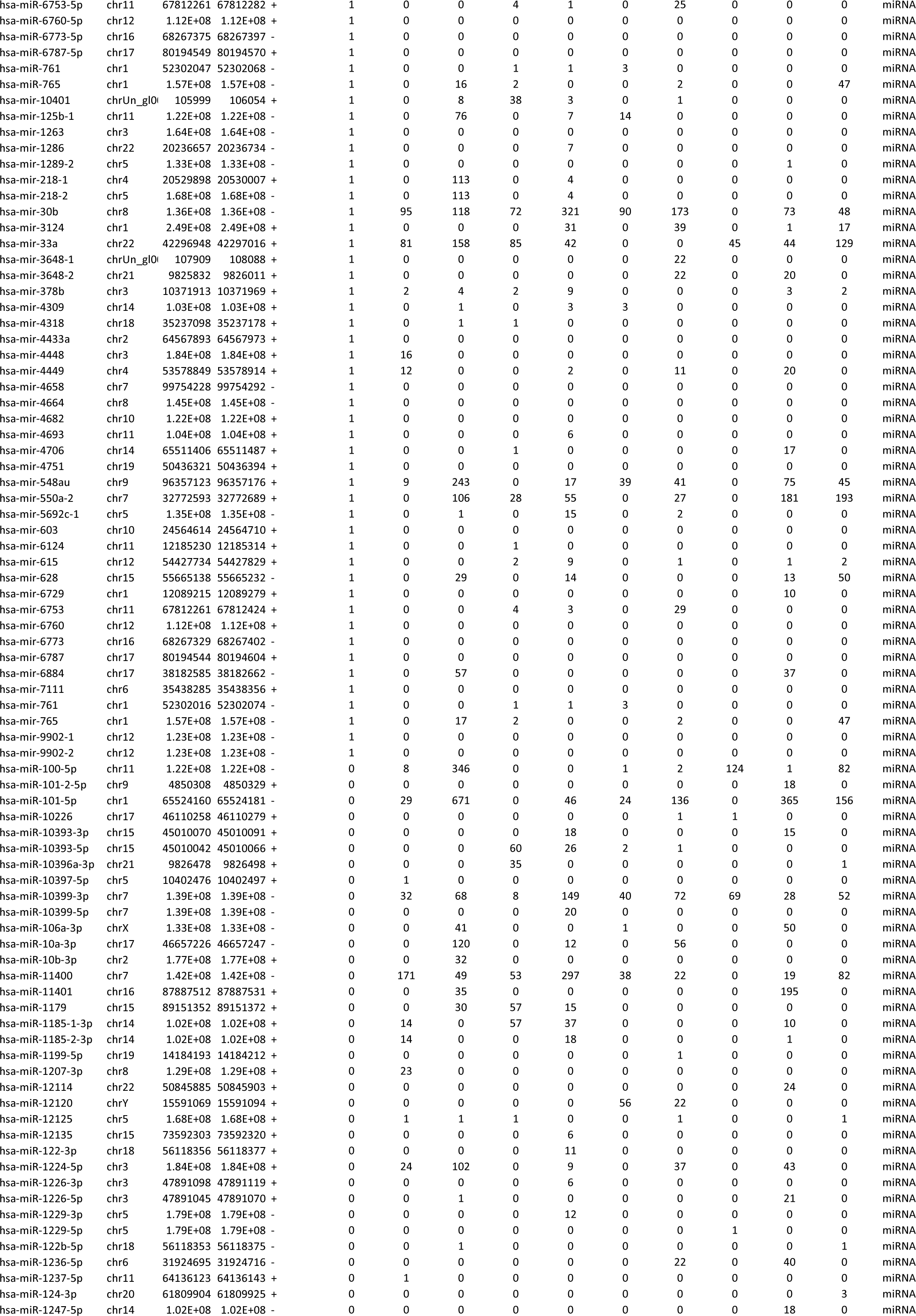
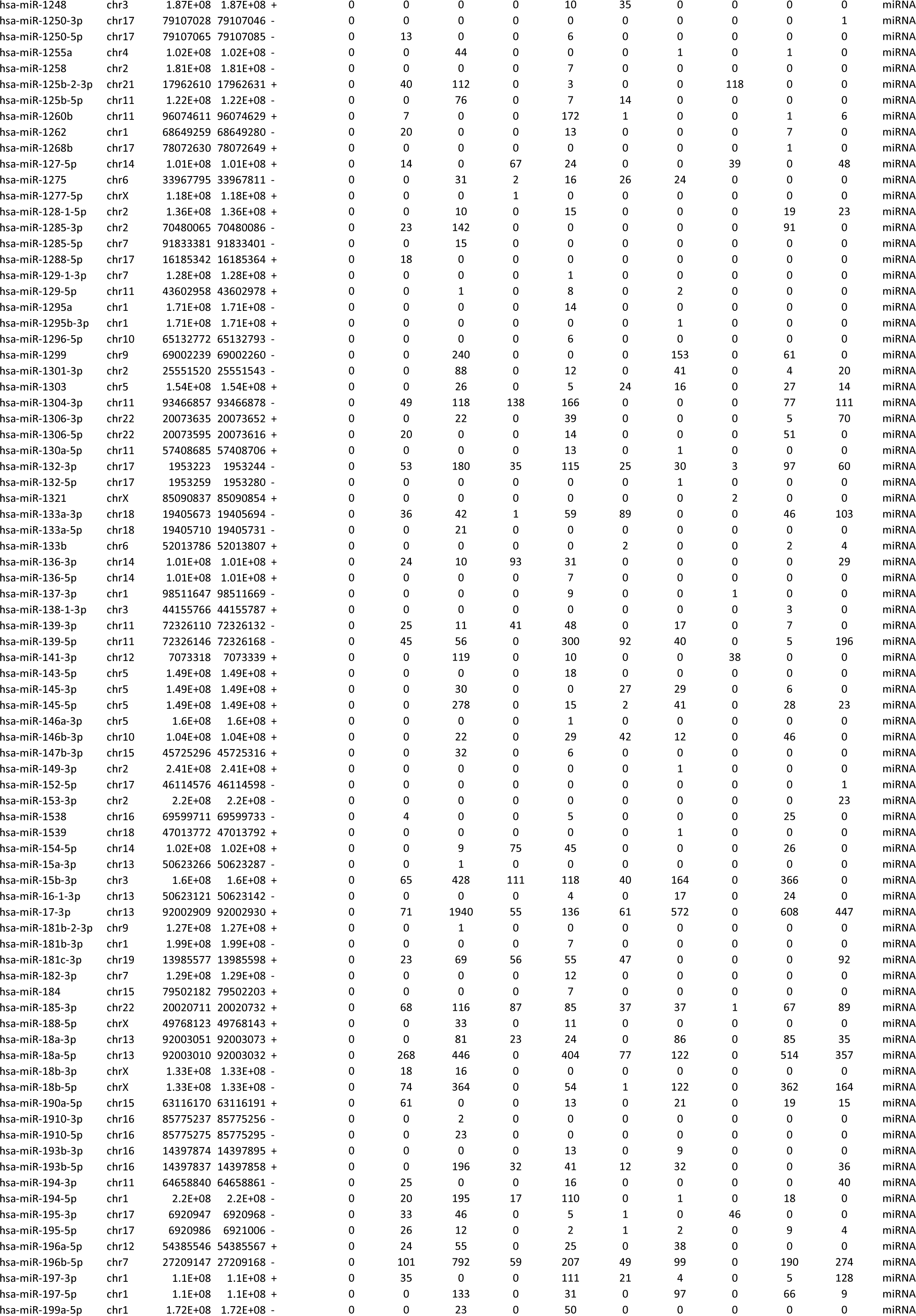
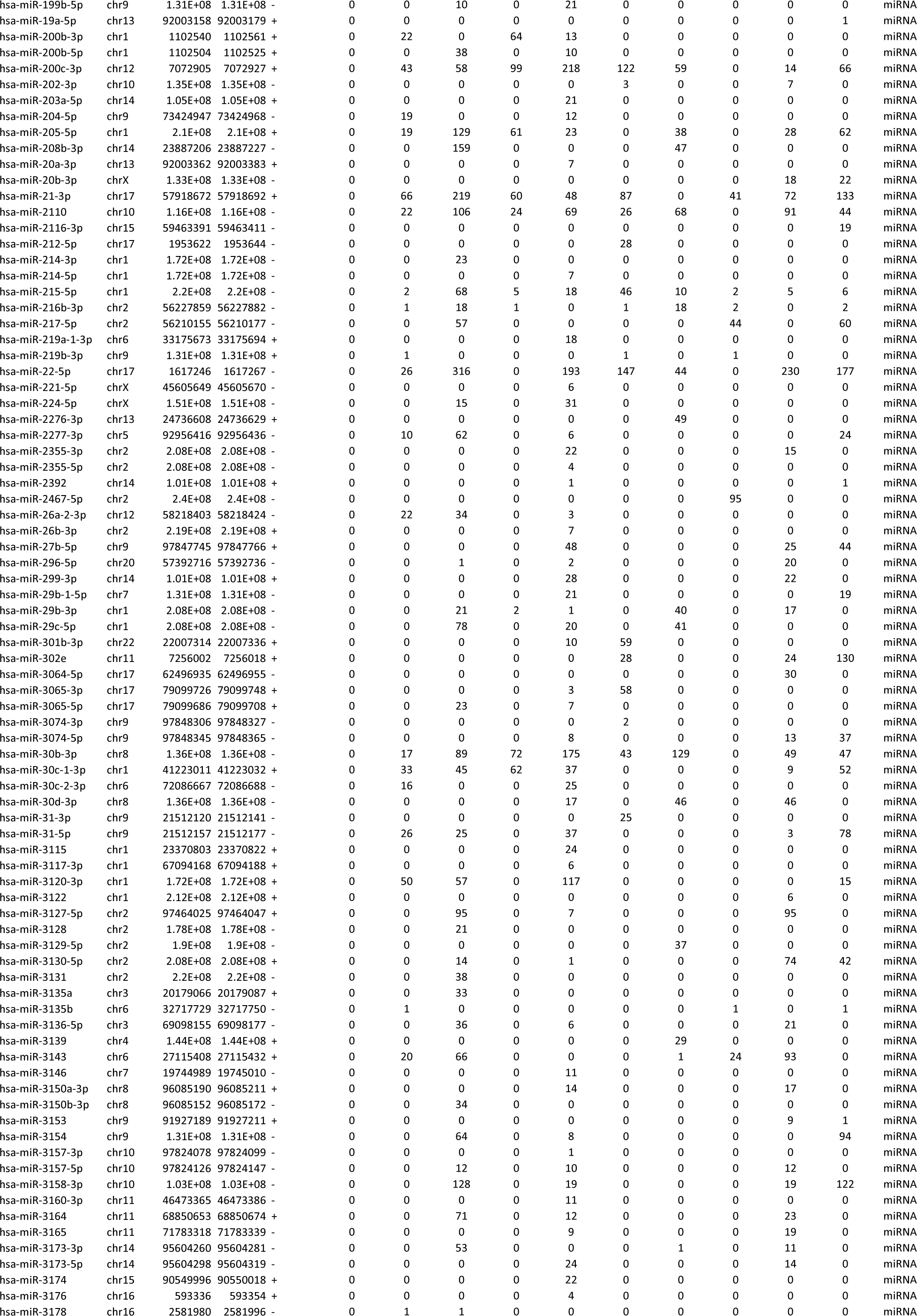
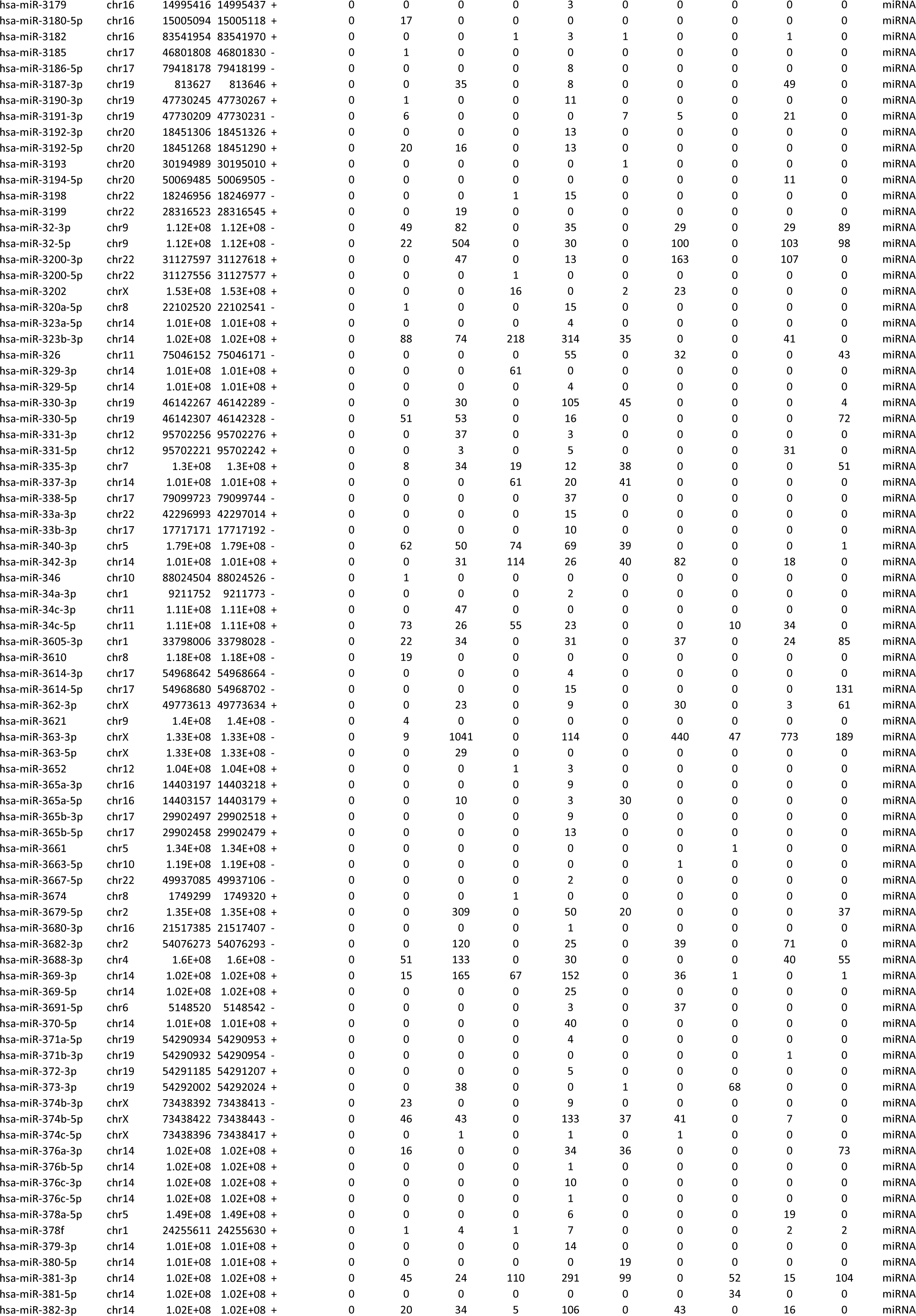
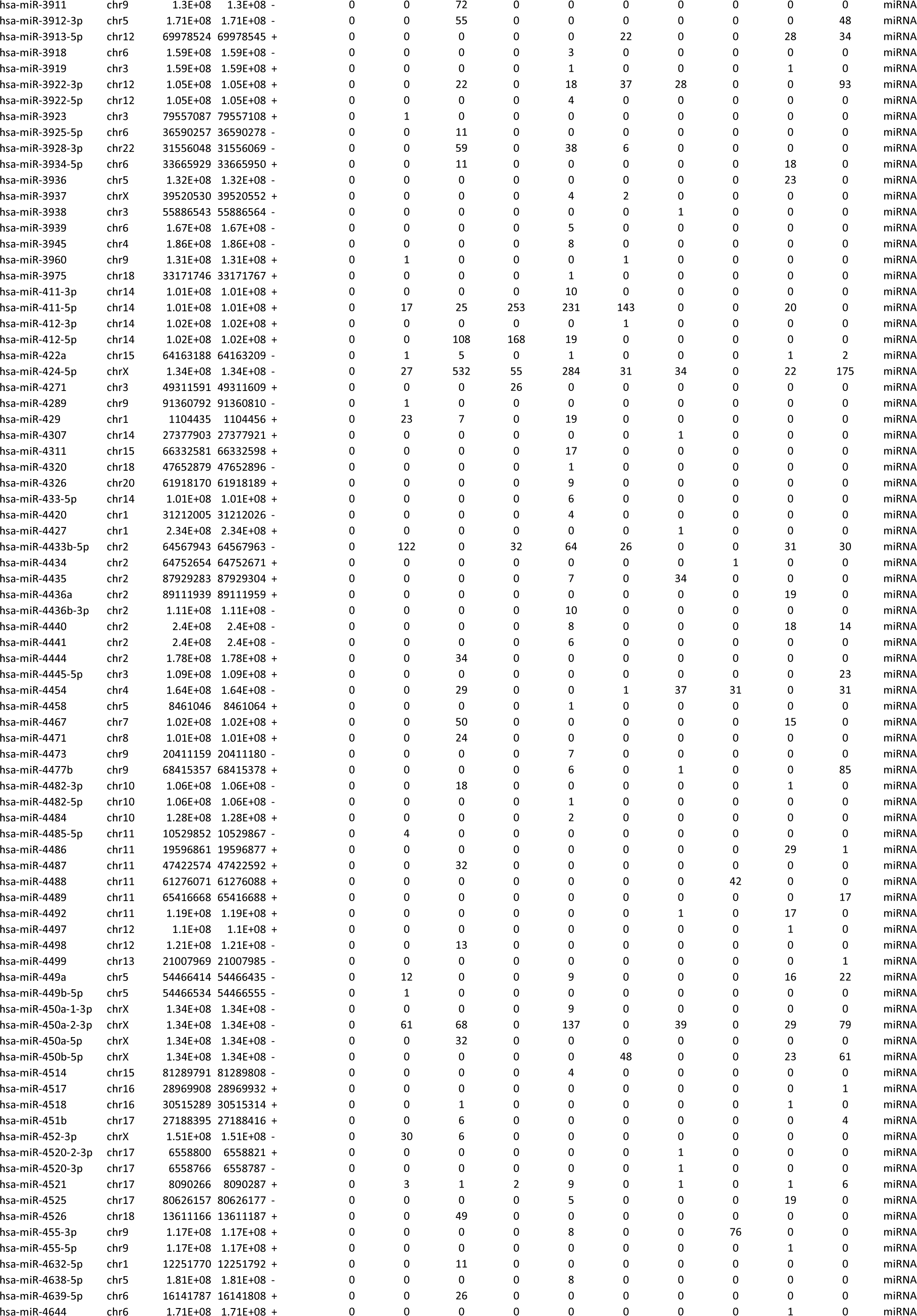
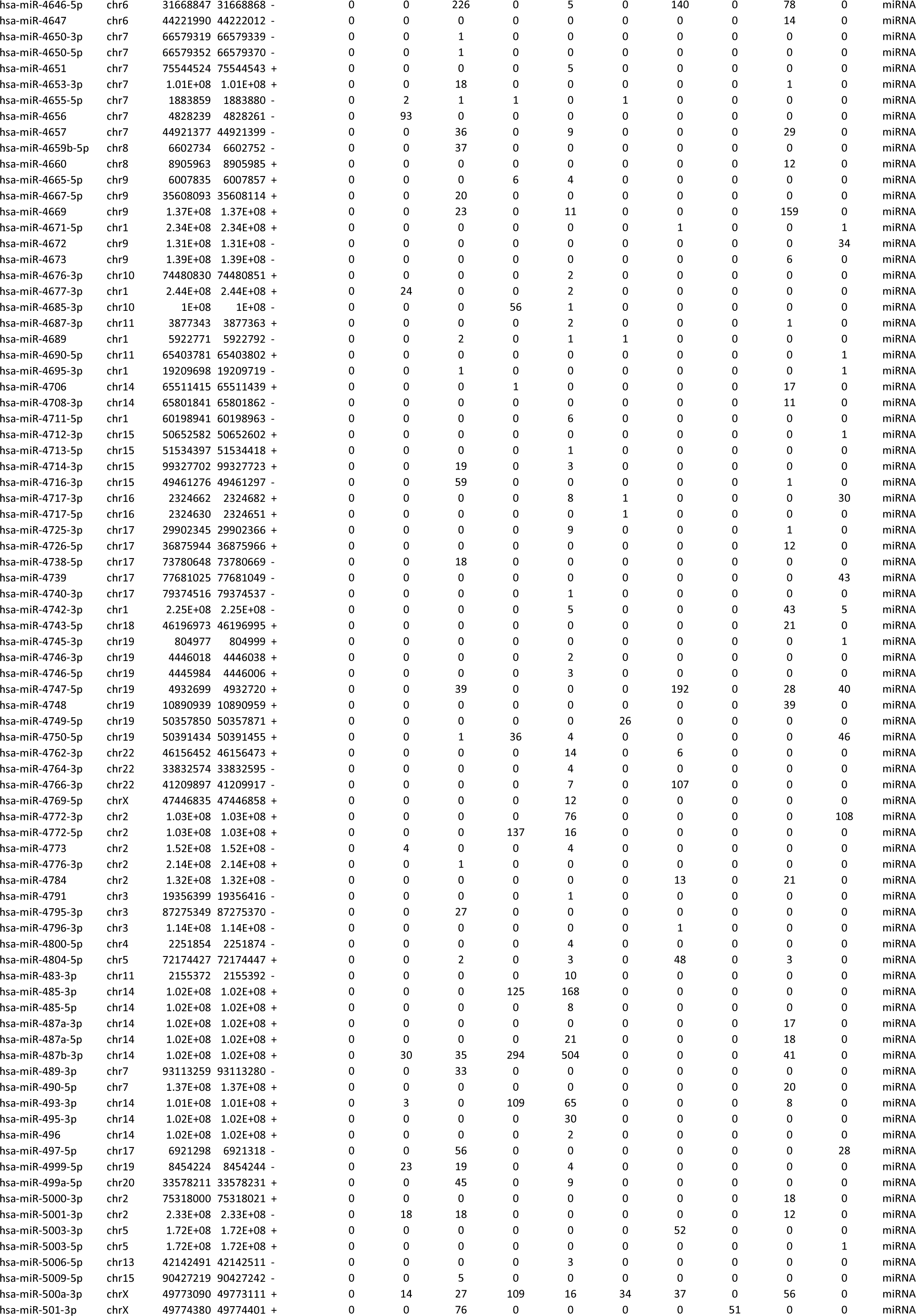
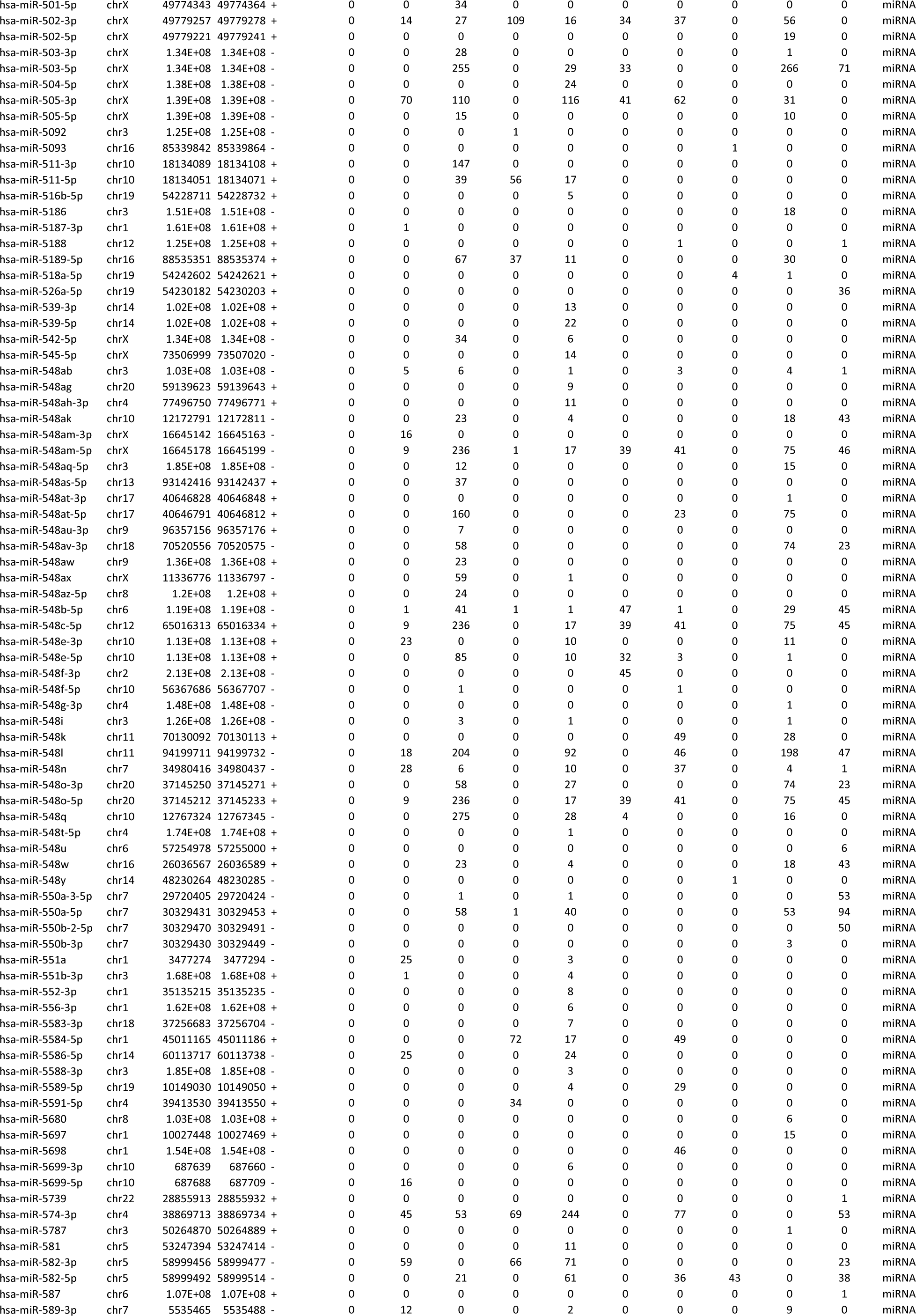
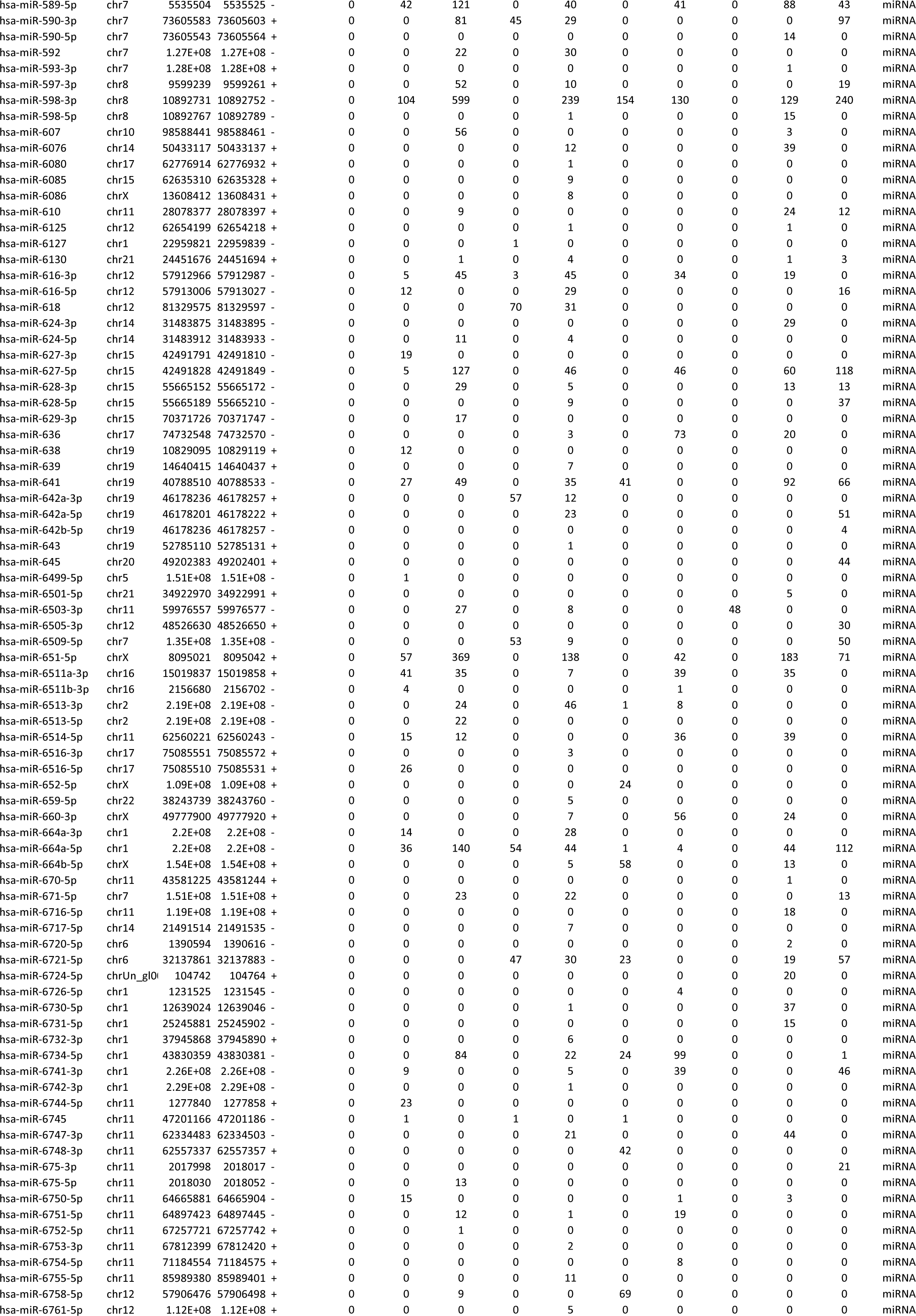
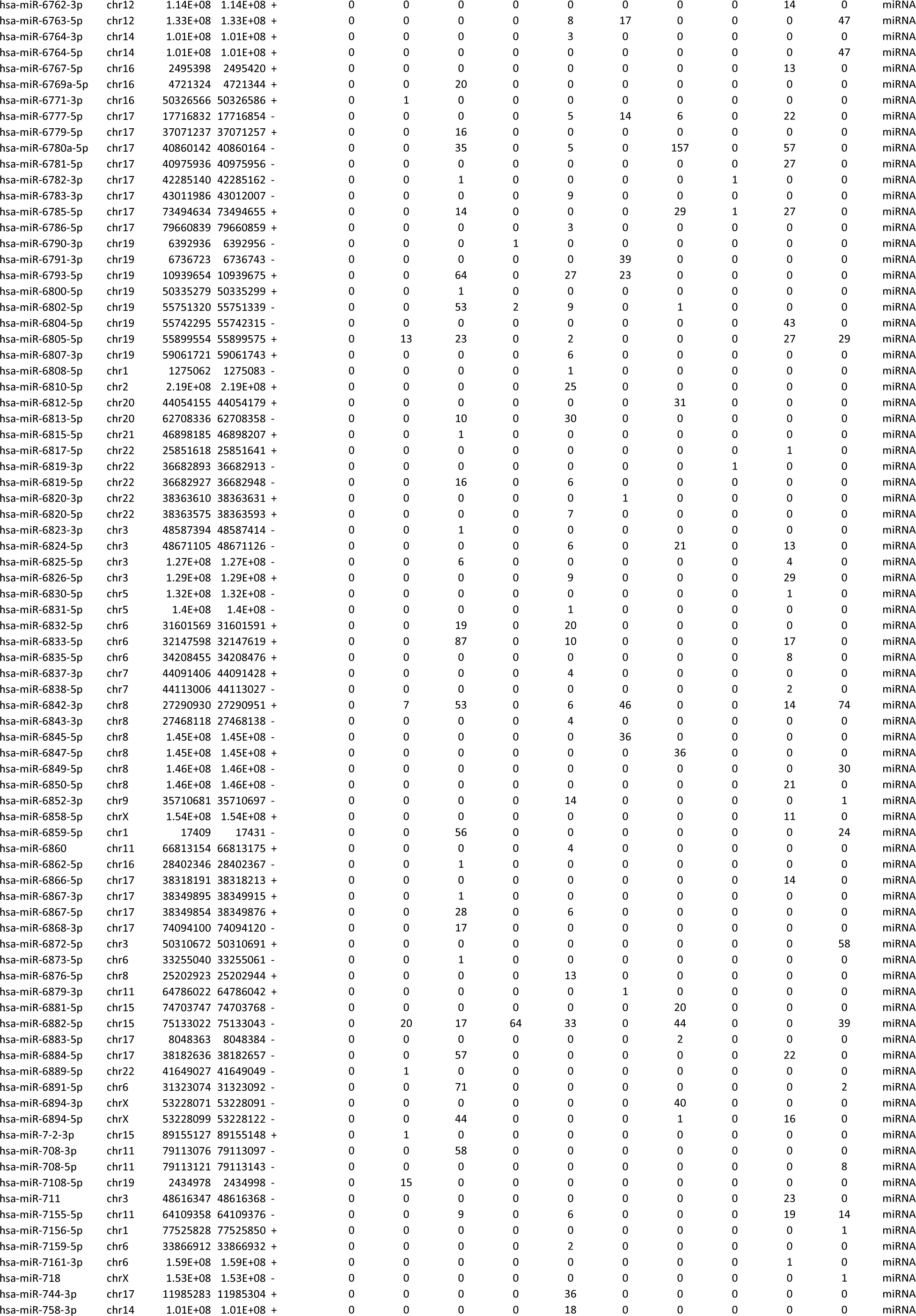
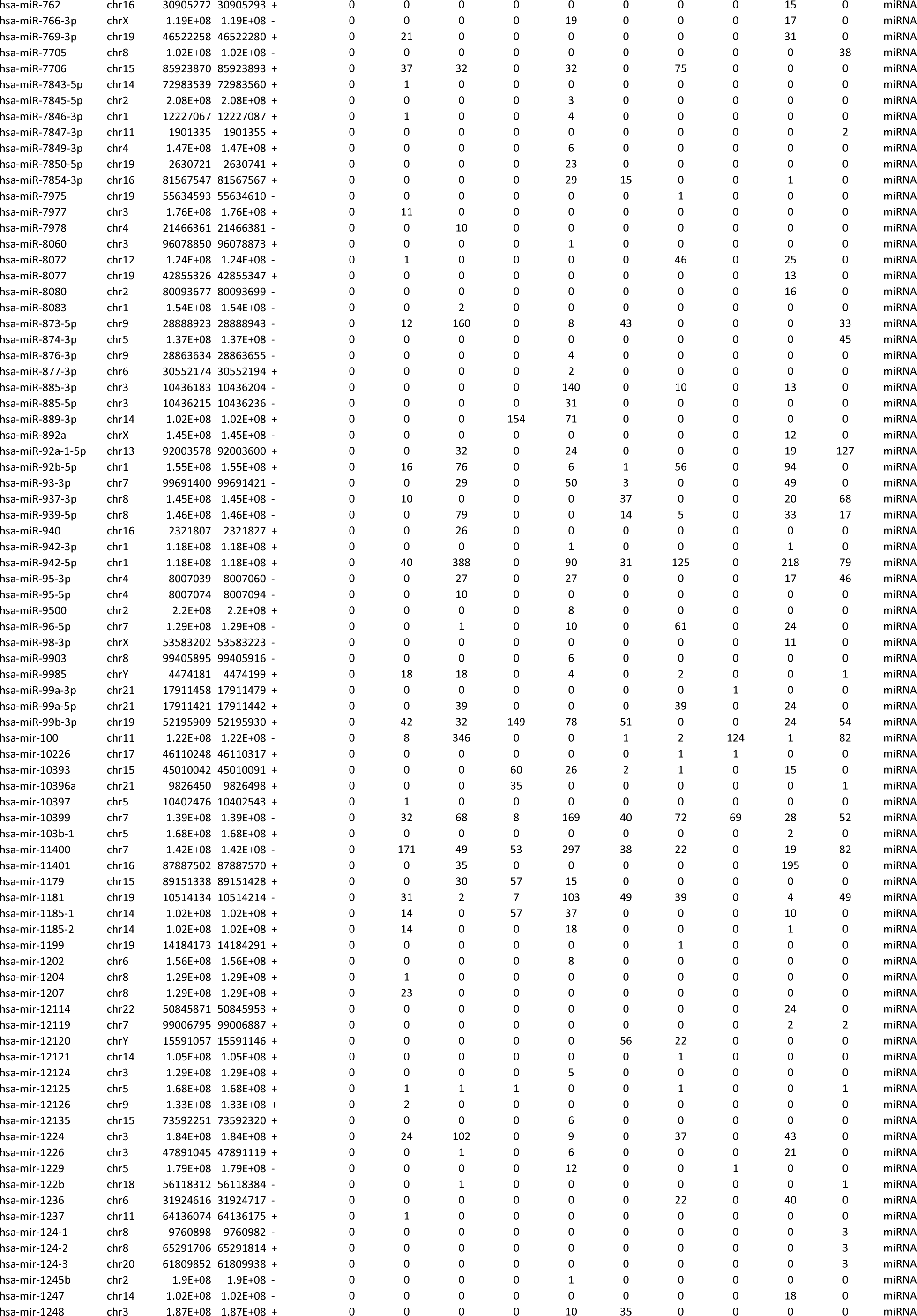
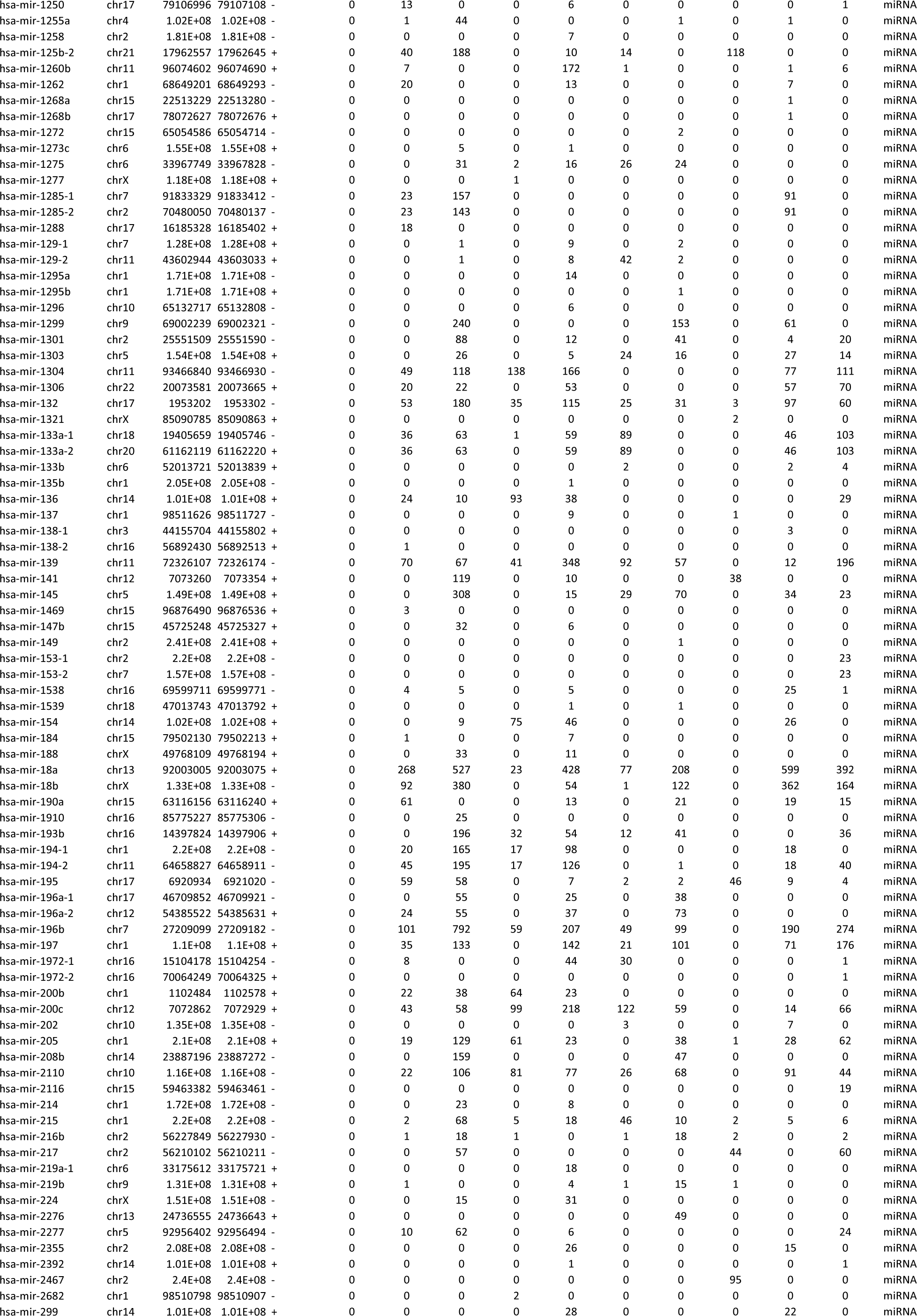
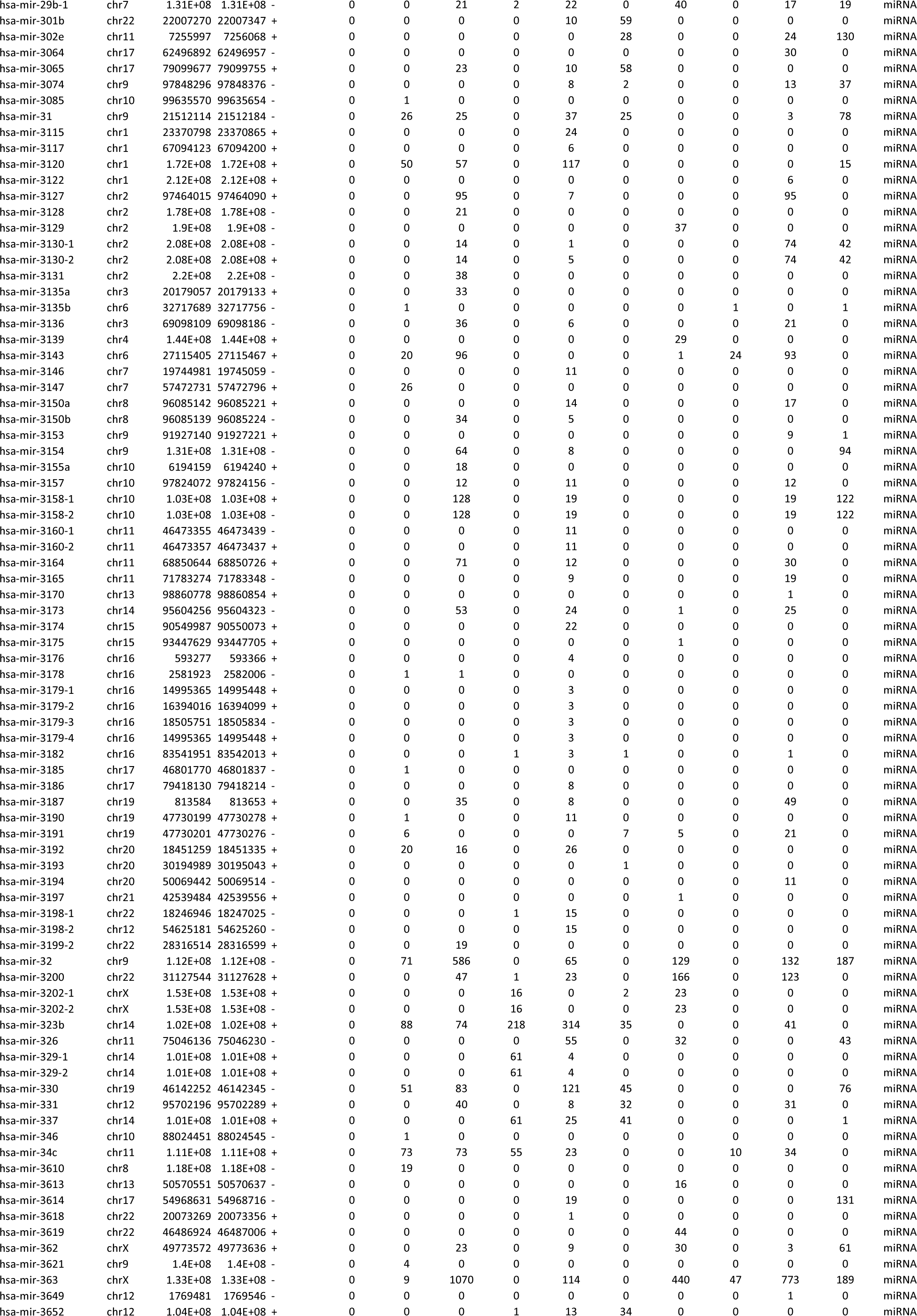
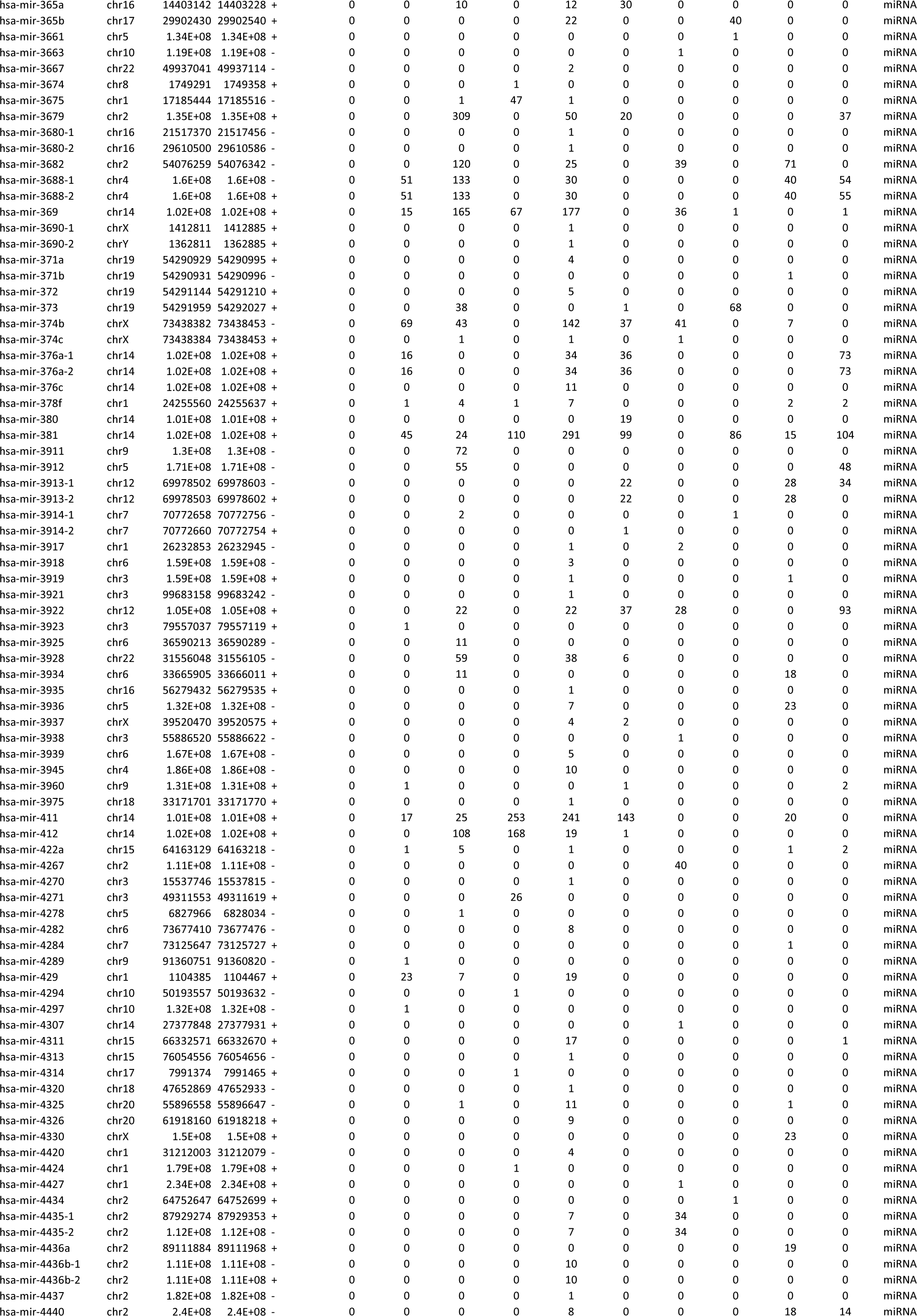
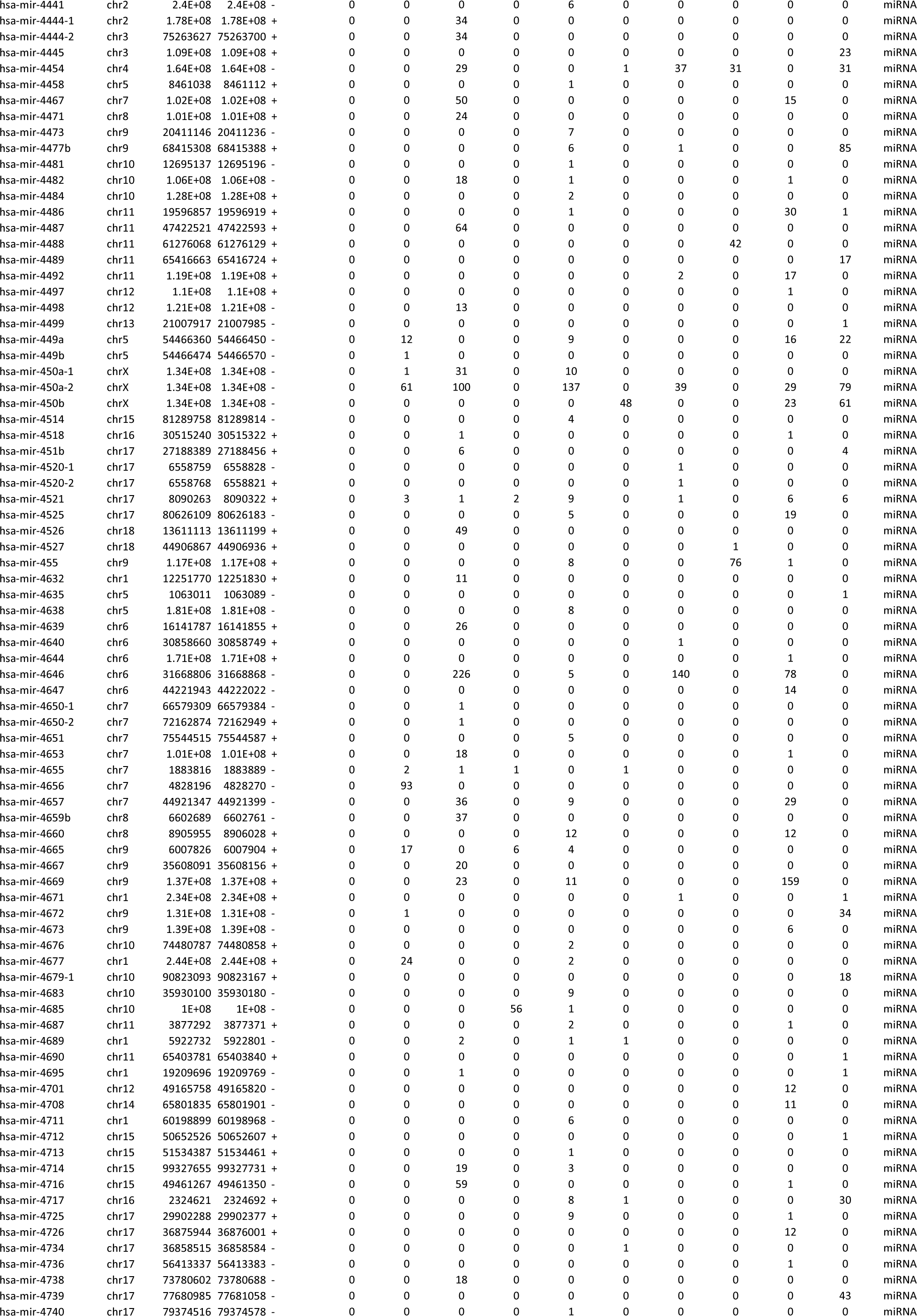
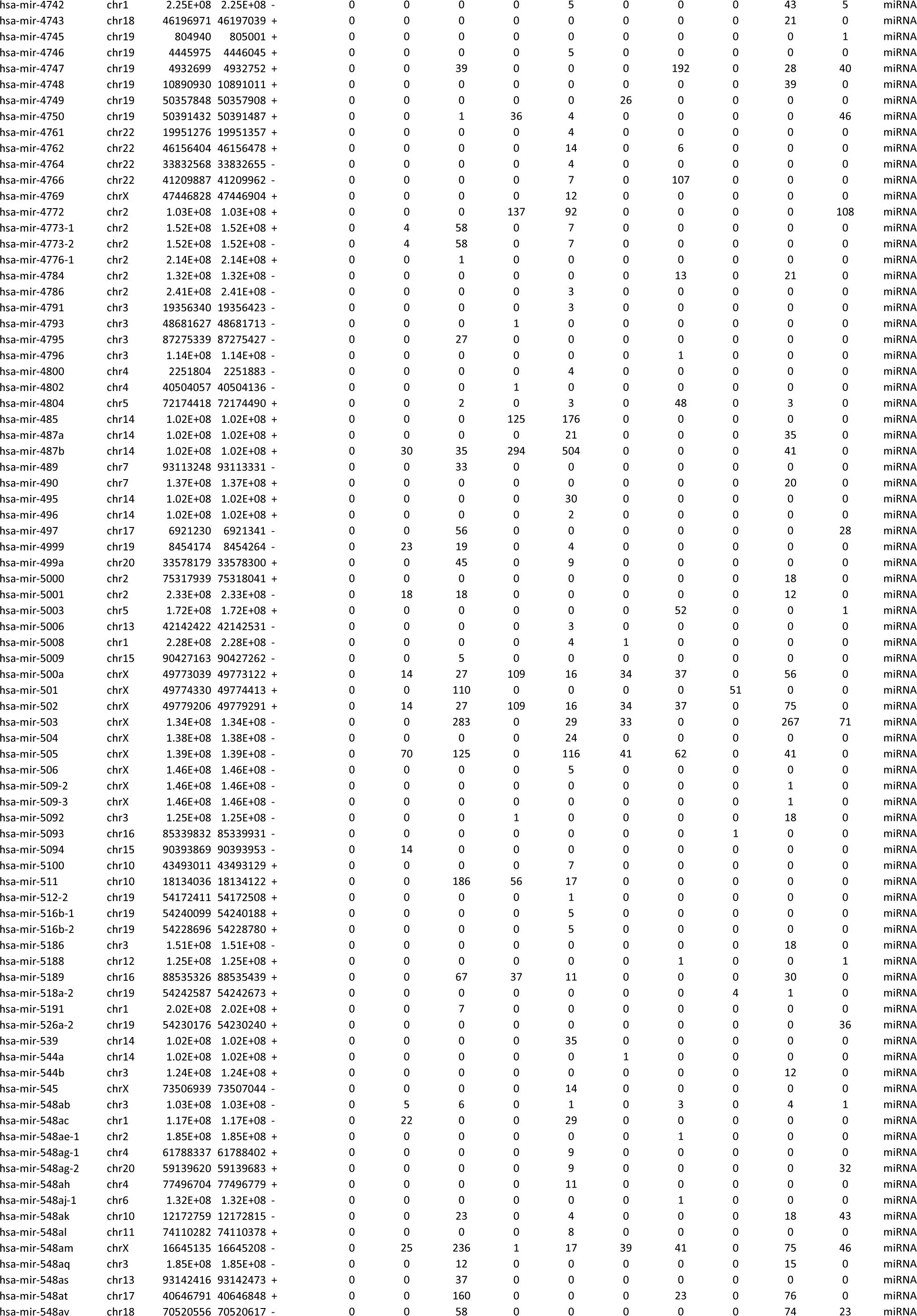
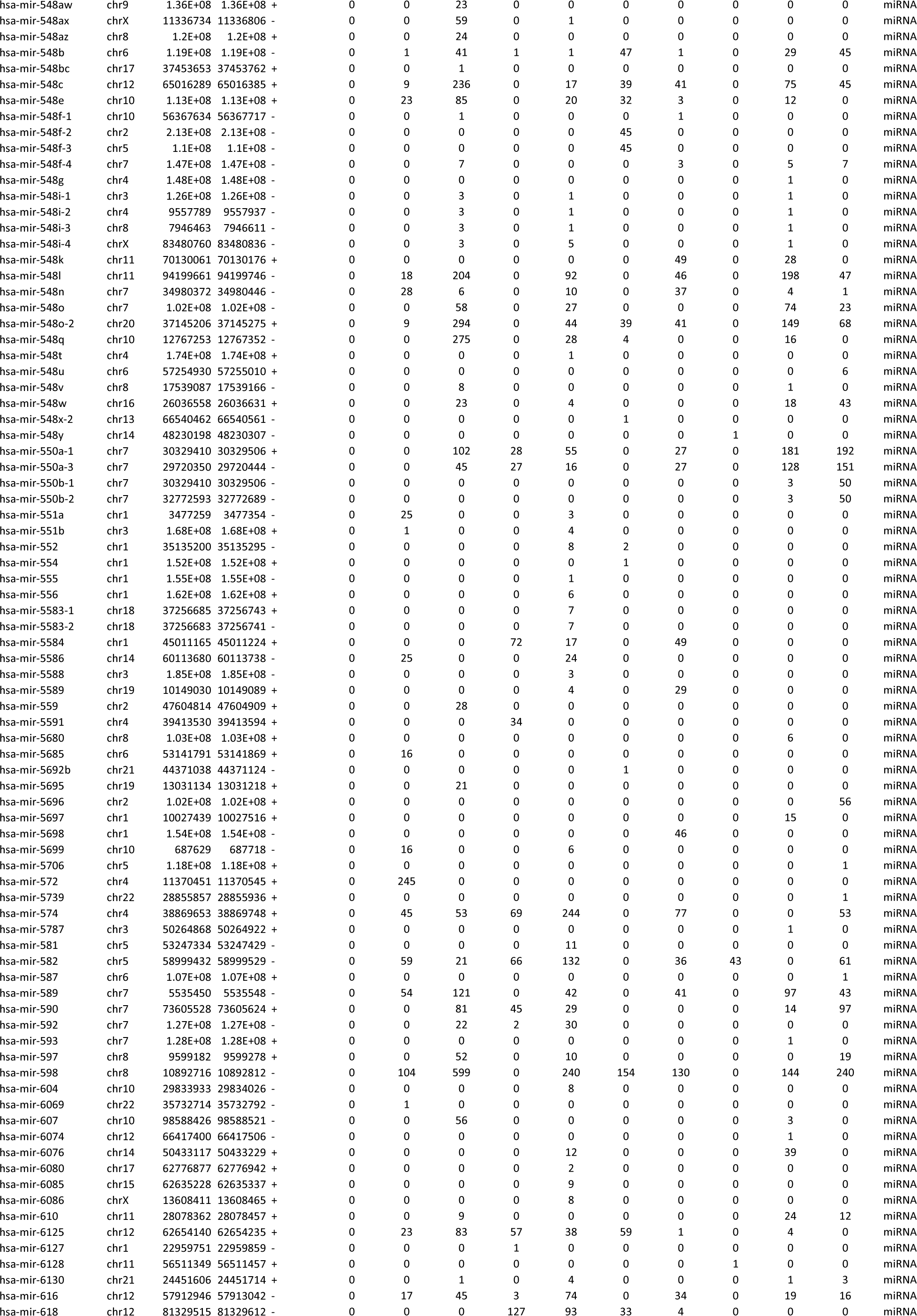
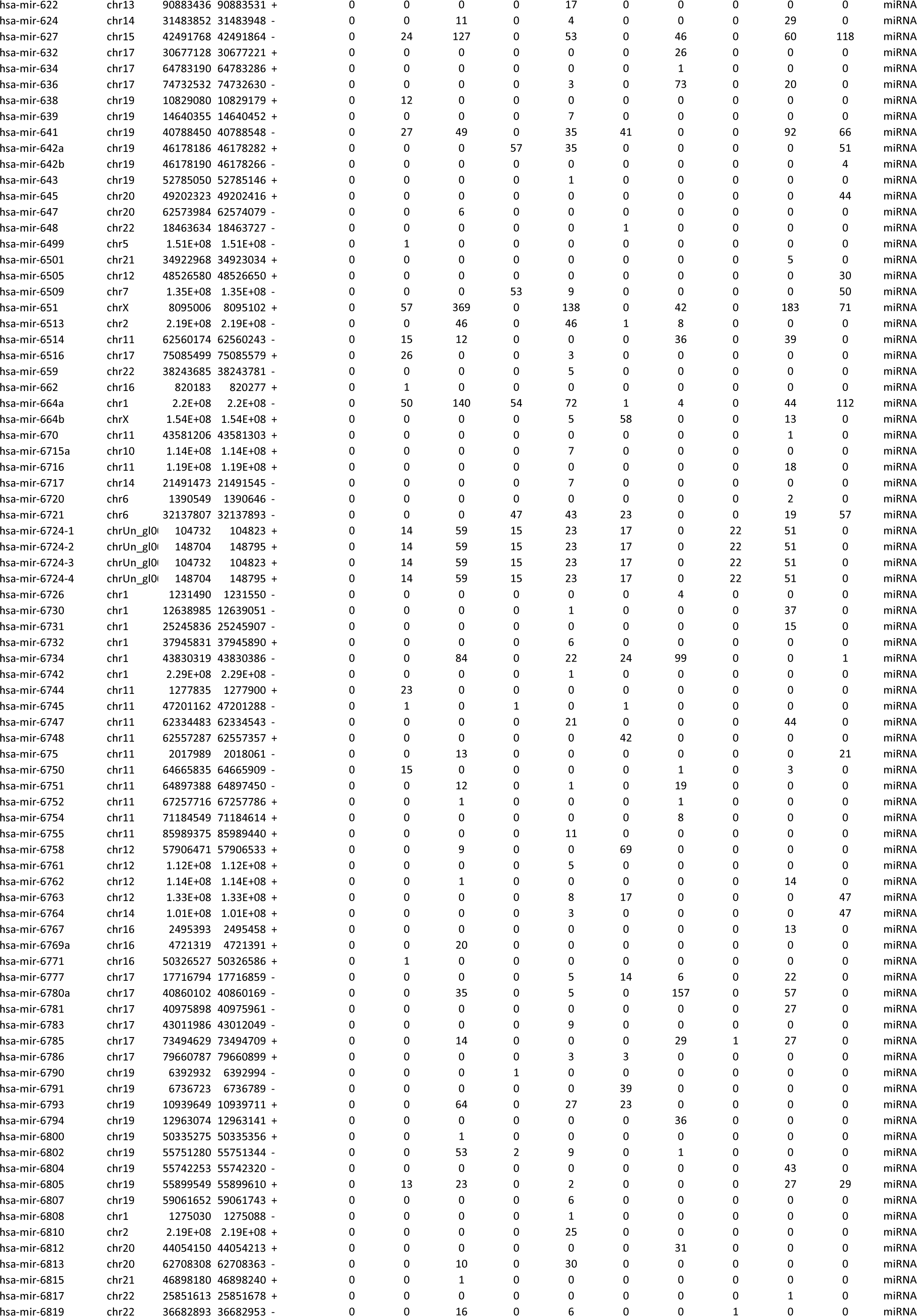
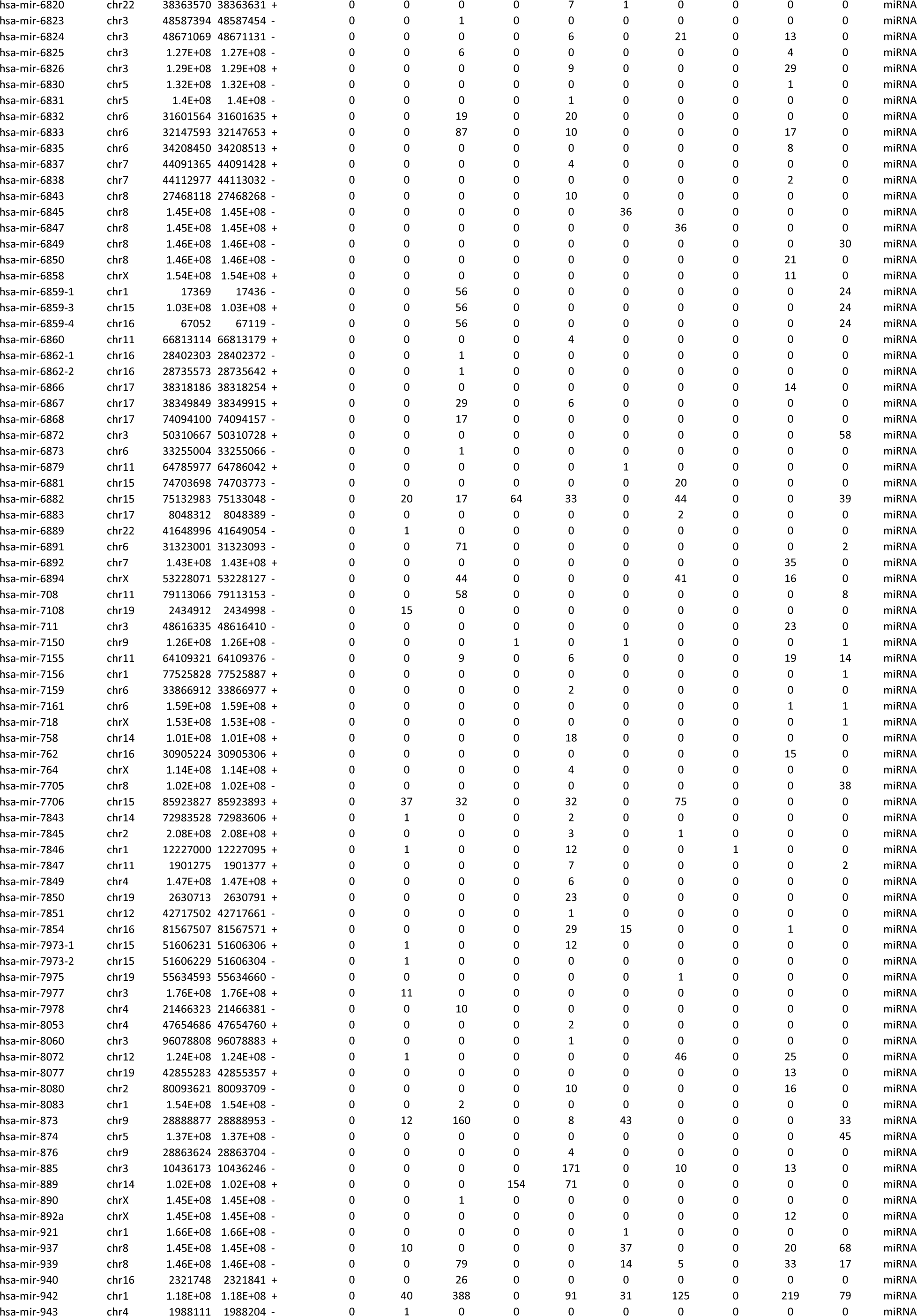
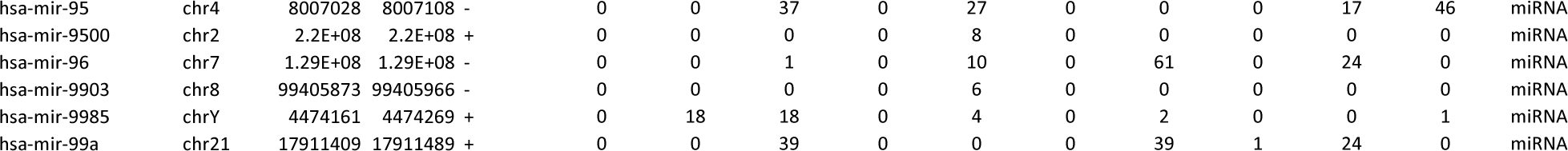

**Supplementary Table-3.** mi-RNAs associated with mitochondrial biogenesis.

| Common Identified mi-RNA in our List vs mi-RNA Data Base |
| --- |
| hsa-miR-16-5p |
| hsa-miR-22-3p |
| hsa-miR-340-5p |
| hsa-miR-15b-5p |
| hsa-miR-182-5p |
| hsa-miR-203a-3p |
| hsa-miR-433-3p |
| hsa-miR-1291-5p |
| hsa-miR-1291 |
| hsa-miR-323a-3p |
| hsa-miR-200a-3p |
| hsa-miR-6876-3p |
| hsa-miR-1185-1-3p |
| hsa-miR-1185-2-3p |
| hsa-miR-132-3p |
| hsa-miR-16-1-3p |
| hsa-miR-181c-3p |
| hsa-miR-195-5p |
| hsa-miR-26b-3p |
| hsa-miR-3120-3p |
| hsa-miR-3146 |
| hsa-miR-3158-3p |
| hsa-miR-34a-3p |
| hsa-miR-362-3p |
| hsa-miR-424-5p |
| hsa-miR-4712-3p |
| hsa-miR-4746-3p |
| hsa-miR-497-5p |
| hsa-miR-5001-3p |
| hsa-miR-503-5p |
| hsa-miR-6762-3p |
| hsa-miR-6813-5p |
| hsa-miR-6826-5p |
| hsa-miR-6838-5p |
| hsa-miR-885-5p |
| hsa-miR-3146 |
| hsa-miR-5696 |

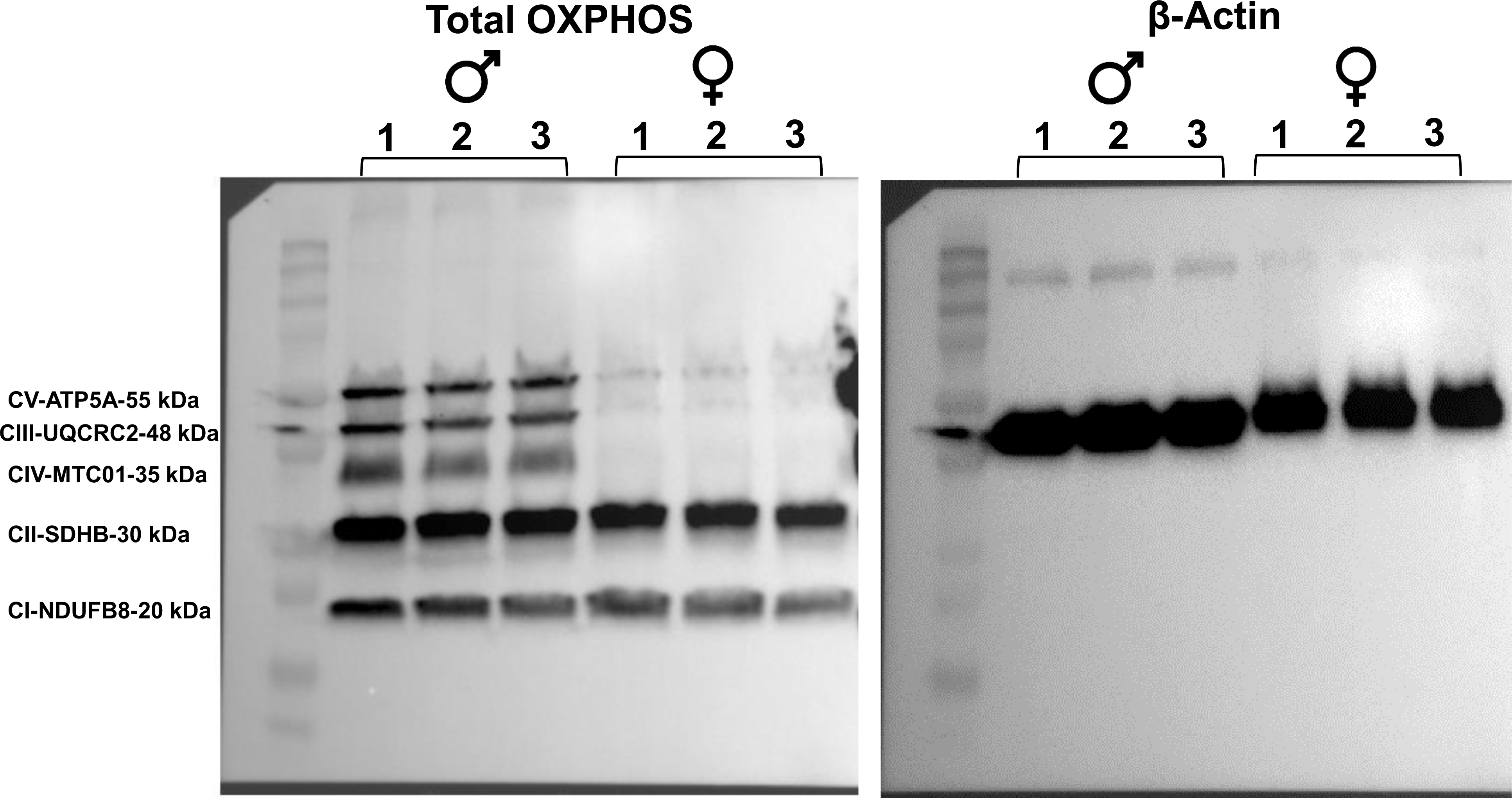

## Notes

### Competing Interest Statement

The authors have declared no competing interest.

